# Structural basis for fork reversal and RAD51 regulation by the SCF ubiquitin ligase complex of F-box helicase 1

**DOI:** 10.1101/2025.09.30.679681

**Authors:** Briana H. Greer, Javier Mendia-Garcia, Elwood A. Mullins, Emma M. Peacock, Sander K. Haigh, Carl J. Schiltz, Clara Aicart-Ramos, Miaw-Sheue Tsai, David Cortez, Fernando Moreno-Herrero, Brandt F. Eichman

## Abstract

Replication fork reversal helps maintain genomic stability during replication stress. F-box helicase 1 (FBH1) catalyzes fork reversal and is an SCF (SKP-CUL1-F-box) E3 ubiquitin ligase that limits RAD51 association with chromatin. Here, we show that preferential binding of SCF^FBH1^ to the lagging strand template at DNA fork structures stimulates helicase activity and is required for fork reversal. A cryo-EM structure of SCF^FBH1^ bound to DNA representing a stalled fork reveals an intimate interaction between FBH1 and the fork junction. Disruption of this interface severely curtails fork reversal *in vitro* and replication progression in cells, providing a model for how ssDNA translocation by FBH1 facilitates annealing of parental DNA by a fundamentally different mechanism than the fork remodelers SMARCAL, HLTF, and ZRANB3. The structure also provides a model for SCF^FBH1^ disassembly of RAD51 filaments through translocation and ubiquitination, and implies that RAD51 is associated with the lagging strand at stalled forks.

## Introduction

Progression of the replisome is challenged by a number of sources of replication stress, including DNA damage, transcription conflicts, DNA repeats and secondary structures, and nucleotide pools^1, 2^. These impediments can stall replication forks, leading to unstable DNA structures that are prone to nucleolytic cleavage and genomic rearrangements. DNA damage tolerance pathways—translesion synthesis, repriming, template switching, and fork reversal—ensure continued replication, thereby maintaining genomic integrity^2, 3, 4^. Dysregulation of these pathways or malfunction of the enzymes involved creates genomic instability and leads to heritable diseases, underscoring their importance.

Replication fork reversal entails the reannealing of parental template strands, unwinding of nascent strands from their templates, and annealing of the two nascent strands^5, 6, 7^. The resulting four-way Holliday junction enables fork restart through recombination, error-free lesion bypass through template switching, and potentially facilitates DNA repair of a fork-stalling lesion by placing it back into the context of dsDNA. Fork reversal is a highly regulated process as prolonged Holliday junctions increase the likelihood of nascent strand degradation or double-strand breaks (DSBs)^8^. Several ATP-dependent DNA translocases are known to catalyze fork reversal *in vivo* and *in vitro*^7, 9^. Loss or dysfunction of fork remodelers leads to genomic instability, increased replication stress, and various diseases, including cancer^9^. HLTF, SMARCAL1, and ZRANB3 are SWI/SNF2-related dsDNA translocases that anneal template strands by binding to the parental duplex, similar to that observed for the bacterial fork reversal enzyme RecG^10, 11, 12, 13, 14^. Whereas the SNF2 remodelers lack DNA unwinding (helicase) activity, several DNA helicases (e.g., BLM, WRN, RECQ5, FANCM)^7, 15, 16, 17^ have been implicated in fork reversal. How DNA unwinding can facilitate reannealing of parental DNA is unknown.

FBH1 is a superfamily (SF) 1 DNA helicase^18, 19, 20^ that catalyzes fork reversal *in vivo* and *in vitro*^21^ and is essential for the replication stress response^22^. Genetic analysis suggests that FBH1 acts in a distinct pathway from SMARCAL1, HLTF, and ZRANB3 based on a unique set of factors required to protect their reversed forks from nucleolytic degradation^23, 24^. FBH1 promotes ATM-mediated signaling in response to DSBs and is required for G2/M-phase checkpoint control following replication stress^25, 26^. Depletion or mutation of FBH1 results in increased sensitivity to DNA damage, unchecked cell cycle progression, and decreased apoptosis^21, 25, 27, 28^. In response to hydroxyurea, FBH1 limits replication fork progression by PRIMPOL^29^. FBH1 also negatively regulates recombination during replication stress and has been shown to process recombination intermediates in mitosis and meiosis^30, 31, 32, 33, 34, 35, 36, 37^. Loss of FBH1 activity is often seen in cancer cells, specifically melanoma and carcinoma^38, 39^. FBH1 protects melanocytes from UV-mediated transformation^25^ and FBH1-deficient tumors may be sensitized to WEE1 inhibition, suggesting FBH1 is required for the efficient induction of the replication stress response after treatment with WEE1 inhibitors^40^.

FBH1 contains a UvrD-like SF1 helicase domain that unwinds DNA with 3′-5′ polarity^18^. ATP hydrolysis (ATPase) activity of the helicase domain is required for fork reversal^21, 23^. A functional helicase is also required for FBH1-dependent promotion of DSBs and apoptosis in response to DNA replication stress^41^. Similar to the UvrD-related, anti-recombinase yeast Srs2^42, 43^, FBH1 physically interacts with and disassembles RAD51 filaments in an ATPase dependent manner^36, 37, 44^. The protein is recruited to stalled forks by direct interaction with proliferating cell nuclear antigen (PCNA) via an N-terminal PCNA-interacting protein (PIP) box and an AlkB homolog 2 PCNA-interacting motif (APIM) located within the helicase domain^38, 45^. Single-molecule studies suggest that FBH1 unwinds the lagging strand at replication forks, based on a specificity for branched structures^44^.

In addition to its role as a DNA helicase, FBH1 is an E3 component of a SKP1-CUL1-F-box protein (SCF) ubiquitin ligase complex that targets RAD51^33, 46, 47^. The SCF^FBH1^ ubiquitin ligase activity is not required for fork reversal^21, 23^ but is important for the DNA damage response, as F-box mutants increase cellular sensitivity to DNA damaging agents^30, 35, 48, 49, 50^ and regulate FBH1 recruitment and function at sites of DNA damage^27, 50^. SCF^FBH1^ ubiquitination of RAD51 reduces RAD51 association with chromatin^27, 33, 36, 37, 49^. Similarly, both helicase and ubiquitin ligase activities limit accumulation of recombination intermediates by *S. pombe* Fbh1^34, 35, 48^. Thus, both ubiquitin ligase and translocation activities of FBH1 prevent unscheduled recombination during replication stress by limiting RAD51’s association with chromatin^27, 33, 36, 49, 51^.

Despite the importance of SCF^FBH1^ to the DNA damage response, the mechanisms of fork reversal and regulation of RAD51 are unclear. FBH1 has been proposed to facilitate fork reversal by unwinding the nascent lagging strand^21^, but the details for how this works have not been investigated. Here, we describe the substrate specificity, DNA translocase and fork reversal activities, and a cryo-EM structure of SCF^FBH1^ bound to a 3-way DNA fork. We found that SCF^FBH1^ preferentially binds replication fork junctions on the lagging strand template and identified an important DNA junction binding surface on FBH1, mutation of which severely impairs fork reversal but has only a modest effect on helicase/translocase activity. We also show that the helicase domain is unique among SF1 helicases and adopts a configuration consistent with its role in ubiquitination of RAD51 and disassembly of RAD51 filaments, providing a basis for FBH1’s anti-recombinogenic properties.

## Results and Discussion

All experiments were carried out with the SCF^FBH1^ complex, composed of FBH1 isoform 4, SKP1, CUL1, and RBX1 (**Supplementary Figure S1**). The helicase activities of FBH1 and SCF^FBH1^ were previously shown to be the same^46^. FBH1 isoform 4 lacks the N-terminal 125 residues present in isoform 1. This region contains the PIP box, a non-canonical PIP degron that regulates CRL4^Cdt2^-dependent ubiquitination and degradation of PCNA-bound FBH1^45^, and 11 putative phosphorylation sites of unknown significance. However, because isoform 1 residues 1-117 are predicted to be disordered and previous biochemical studies used isoform 4^18, 36, 46^, we chose isoform 4 for our biochemical and structural studies. All residue numbers reported herein refer to isoform 4 (**Supplementary Fig. S1**).

### FBH1 helicase activity is stimulated at forks

A previous study showed FBH1 to exhibit longer dwell times and repetitive shuttling on a DNA fork substrate than on a duplex containing a 3′-overhang, suggesting that FBH1 has a preference for fork junctions^44^. To test this, we compared DNA binding and unwinding activities of purified SCF^FBH1^ on fork and duplex substrates (**Fig. 1, Supplementary Fig. S1**). Modeling experiments based on other SF1 helicases and preliminary helicase experiments with FBH1 indicated that 5-8 nucleotides of ssDNA are sufficient for optimal activity. Therefore, electrophoretic mobility shift assays were used to measure SCF^FBH1^ binding to a DNA duplex containing an 8-nucleotide 3′-overhang and to 3-way forks containing 8-nucleotide ssDNA regions (gaps) on what would be leading and lagging template strands (**Fig. 1a**). The protein complex exhibited the highest affinity for the fork containing a gap on the lagging strand, and weaker affinity for the leading gap fork and the fork with no gaps. Interestingly, multiple SCF^FBH1^-associated bands were observed for all three forks, suggesting that at higher protein concentrations more than one protein is able to bind. The weakest affinity was observed for the 3′-overhang, consistent with dwell times from the previous study^44^.

**Figure 1.**
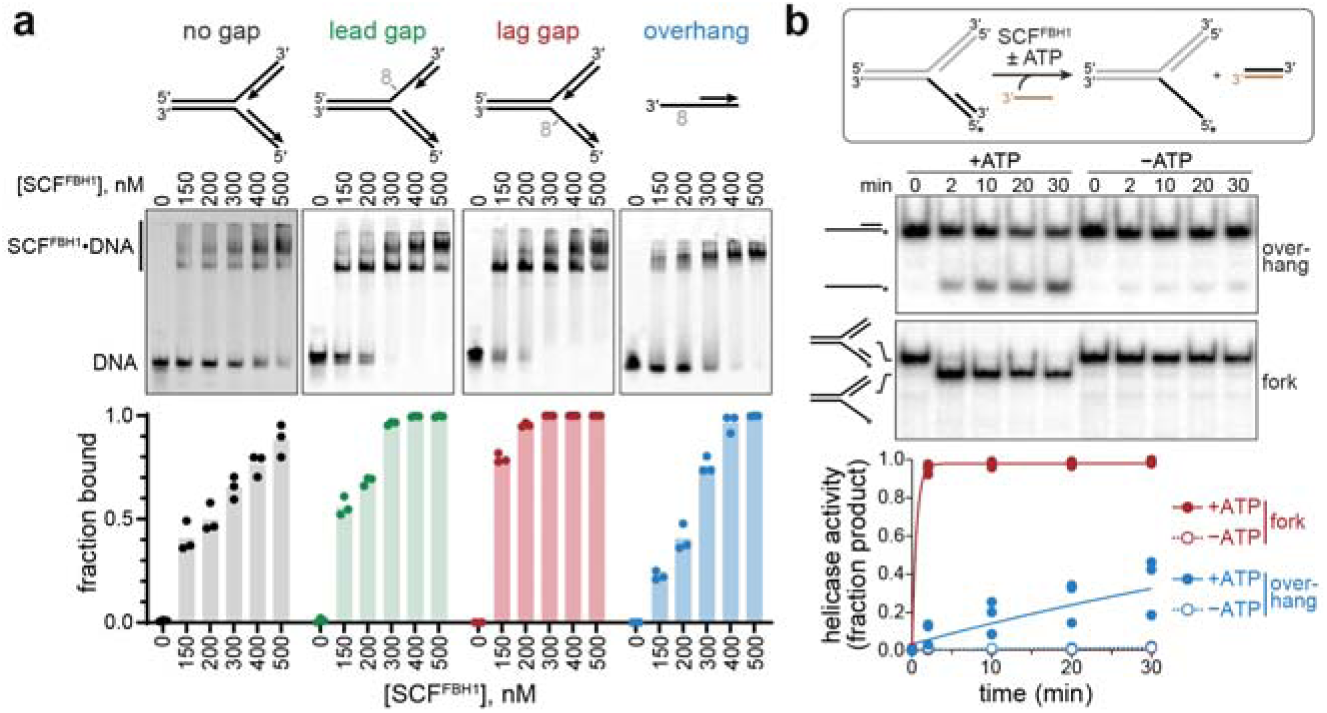
SCF^FBH1^ helicase activity is stimulated at forks. **a**. EMSA data for SCF^FBH1^ binding to for and overhang structures. Representative gels are shown above quantified data. SCF^FBH1^•DNA complex bands used in quantification is defined by the bar to the left of the gels. Bars denote average value (n=3). **b**. Helicase activity. *Top*, schematic of the strand displacement assay, in which the black strand are identical between overhang and fork substrates. The asterisk denotes the position of a FAM label. *Middle*, representative native PAGE separation of overhang and fork substrates and products at different reaction times. *Bottom*, quantification of multiple experiments (n=3-5), fit to a single exponential.

Helicase activity was measured using a strand displacement assay with fork and duplex/3′-overhang substrates that contained 9-nucleotide ssDNA binding regions (**Fig. 1b**). The translocating strand contained a 5′-FAM label for visualization of substrate and products after separation on native polyacrylamide gels, and reannealing of the displaced strand was prevented by adding an excess of unlabeled complementary strand to the reaction. Consistent with the binding data, the rate of strand displacement was at least two orders of magnitude greater for the fork relative to the overhang substrate, with observed rate constants (*k*_obs_) of >1.8 ± 0.1 min^−1^ (fork) and (1.2 ± 0.3)×10^−2^ min^−1^ (overhang) (**Fig. 1b**). The *k*_obs_ for the fork substrate is a lower limit since the reaction was complete at the first time point. The helicase activities were ATP-dependent as no activity was observed in reactions lacking ATP. Together, these results indicate that SCF^FBH1^ prefers to bind to the lagging strand template immediately adjacent to the fork junction, and that this binding mode stimulates the translocase activity of FBH1.

### SCF^FBH1^ force-dependent unwinding and processive ssDNA translocation

We investigated the helicase and ssDNA translocase rates of SCF^FBH1^ using a single-molecule magnetic tweezers (MT) assay, in which a palindromic ssDNA capable of forming a hairpin is tethered between a magnetic bead and a glass surface (**Fig. 2a**)^52^. A constant force below 15 pN that is not strong enough to mechanically unwind the duplex is applied to the bead such that in the absence of helicase activity, the hairpin remains closed. Any helicase-mediated unwinding of the hairpin is detected as an increase in extension of the molecule. Once the helicase completely unwinds the hairpin, it continues translocating on the ssDNA and the duplex reforms in its wake. Thus, ssDNA translocation is measured as a decrease in extension after initial unwinding.

**Figure 2.**
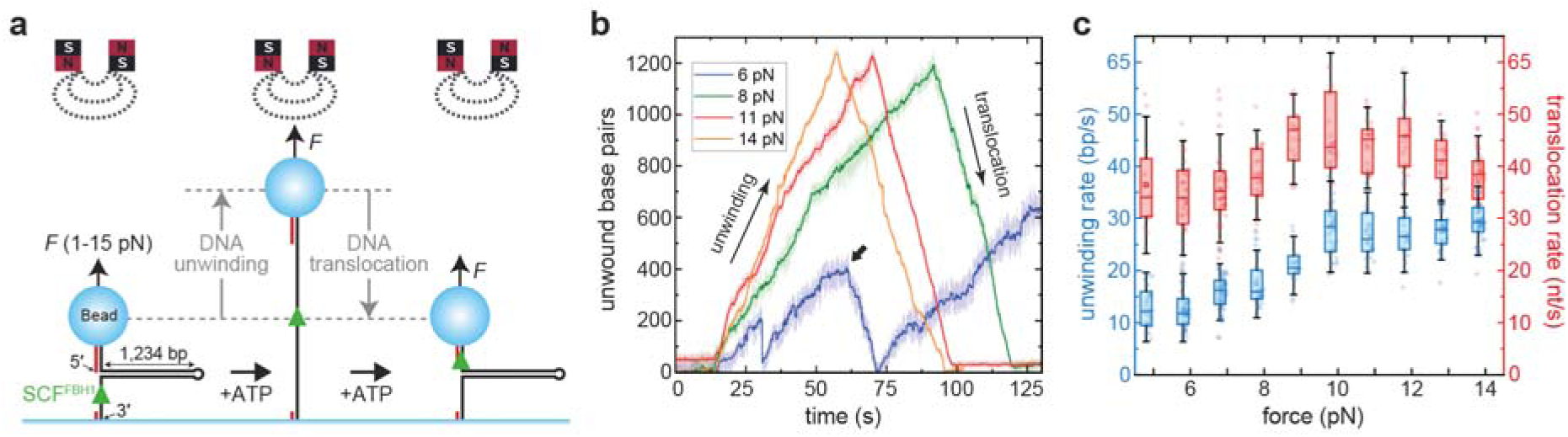
SCF^FBH1^ displays 3′-5′ force-dependent unwinding activity and efficient ssDNA translocation. **a**. Magnetic tweezers hairpin unwinding experimental setup. **b**. Representative time courses of individual activities of SCF^FBH1^ at different forces. The thick arrow in the blue trace indicates a strand switching event. **c**. Quantification of duplex unwinding (blue) and ssDNA translocation (red) velocities measured at different forces applied to the hairpin.

To observe the activity of individual proteins, we measured unwinding and translocation rates at forces ranging from 5–14 pN (**Fig. 2b,c, Supplementary Fig. S2**) at low (100 pM) protein concentration. No unwinding activity was observed below 5 pN, even under conditions designed to trap SCF^FBH1^ between duplex regions by transiently opening and re-closing the hairpin. Above this threshold, SCF^FBH1^ unwound the hairpin with rates that increased modestly with force, from 13 bp/s at 5 pN and plateauing at 28 bp/s at 10 pN and above. Translocation rates were higher than unwinding rates and peaked at 10 pN, where SCF^FBH1^ moved at approximately 46 nt/s, compared to 34 nt/s at 5 pN (**Fig. 2c**). At low forces (5-7 pN), we observed strand switching events, which in our traces are reflected as changes from unwinding to reannealing before the protein reaches the hairpin loop (see arrow in blue trace in **Fig. 2b**). Reannealing events that followed strand switching occurred at 36 bp/s, similar to the enzyme’s translocation rate and more than two orders of magnitude slower than spontaneous hairpin closure measured after protein detachment (**Supplementary Fig. S3**). Thus, the reannealing observed during strand switching reflects active enzyme translocation rather than spontaneous duplex closure. Together, these data indicate that SCF^FBH1^ is a robust helicase/translocase capable of translocating over 2 kb without dissociating, comparable to other SF1 enzymes^52, 53^.

In contrast to what has been observed with SNF2-family remodelers using a similar experimental setup^10, 54^, SCF^FBH1^ lacked annealing activity in this assay. To test for annealing with this substrate, we held the hairpin partially open by applying a force just below its mechanical melting threshold. We confirmed that HLTF and RecG can close the hairpin even under high (14 pN) forces by translocating on dsDNA, thereby annealing the complementary strands without strand-separation activity (**Supplementary Fig. S4**)^10, 54, 55, 56^. In contrast, SCF^FBH1^ exhibited processive unwinding but no reannealing activity (**Supplementary Fig. S4**), suggesting that it uses a distinct mechanism of fork reversal compared to SMARCAL1, HLTF, and ZRANB3.

### FBH1 reverses forks by translocating on the lagging strand

SMARCAL1, HLTF, and ZRANB3 reverse forks by translocating on the parental dsDNA upstream of and toward the fork, using enzyme-specific fork recognition domains to position the ATPase motor domains in a manner observed with bacterial RecG^10, 13, 14^. In contrast, the substrate specificity of SCF^FBH1^ (Fig. 1) suggests that it facilitates fork reversal by translocating 3′-5′ on the lagging ssDNA template with the fork junction behind it^21^. To further test this mechanism, we examined the requirements of fork reversal of SCF^FBH1^ on model fork substrates using a gel-based assay that measures conversion of a three-way fork substrate with complementary arms to a duplex product (**Fig. 3**). Consistent with its specificity as a ssDNA translocase, SCF^FBH1^ was unable to reverse a fork structure that lacked ssDNA gaps (**Fig. 3a**). In contrast, this fork is a substrate for SMARCAL1, HLTF, and RecG^12, 13, 57, 58^ and we confirmed HLTF activity under our assay conditions (**Fig. 3a**). Similarly, we did not detect any SCF^FBH1^ fork reversal activity in a single-molecule magnetic tweezers assay with a fully base-paired 3-way fork substrate like those previously used to study the activities of SMARCAL1, RecG, and UvsW (**Supplementary Fig. S4**)^10, 54, 59^, and we confirmed that HLTF and RecG are active in our single-molecule setup (**Supplementary Fig. S4**). The lack of SCF^FBH1^ activity for a fully base-paired substrate reflects an inability to engage the fork in the absence of ssDNA. Consistent with this, we observed robust activity on a fork containing a lagging strand gap (**Fig. 3b**). We did not, however, observe reversal of a fork with a leading strand gap, indicating that 3′-5′ translocation on the lagging strand is necessary for reversal. A modest amount of parental duplex unwinding was observed as the enzyme moves into the fork from the lead gap configuration (**Fig. 3b**, bottom of gel). We also found that displacement of the nascent lagging strand is not required for reversal, as the enzyme was active on a 5′-flap substrate (**Fig. 3c**). SCF^FBH1^’s fork reversal activity is ATP-dependent; we did not observe activity in the absence of ATP or from an FBH1 Walker B (D573A E574A) mutant (**Fig. 3b,c**). These data are consistent with a model in which FBH1 facilitates parental strand annealing by binding to ssDNA on the lagging strand and translocating 3′-5′ with the junction in its wake.

**Figure 3.**
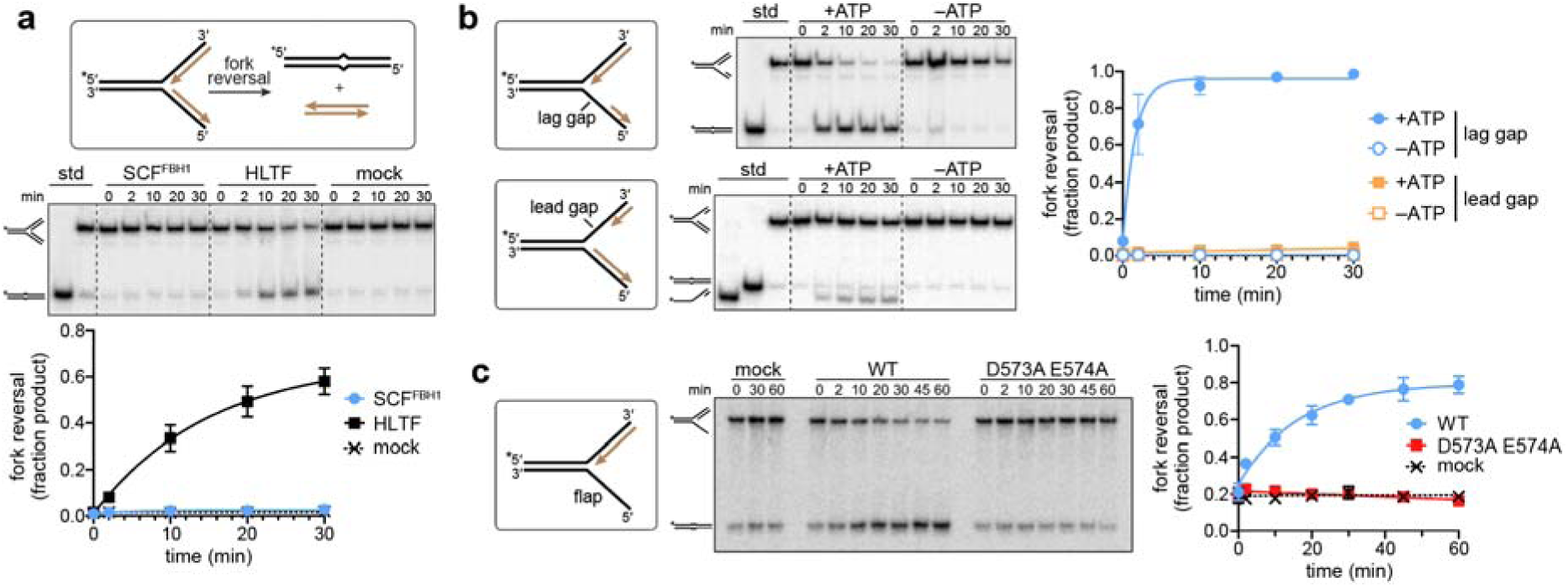
FBH1 reverses forks by translocating on the lagging strand. Fork reversal activity of SCF^FBH1^ on model fork substrates. Each panel shows a representative native PAGE separation of substrate and products and a plot of quantified data (mean ± SD, n=3 or 4). **a**. SCF^FBH1^, HLTF, and no-enzyme (mock) activity on a fork containing no ssDNA gaps. **b**. SCF^FBH1^ activity on forks containing ssDNA gaps on either the lagging (top) or leading (bottom) strands. **c**. Wild-type (WT) and ATPase-deficient Walker B (D573A E574A) SCF^FBH1^ activity on a fork containing no nascent lagging strand (flap substrate).

### Structure of an SCF^FBH1^-ssDNA complex

To further understand the fork reversal and ubiquitin ligase activities of SCF^FBH1^, we determined a cryo-EM structure of the SCF^FBH1^ complex bound to a 3-way fork containing an 8-nucleotide lagging strand gap (**Supplementary Table S1, Supplementary Fig. S5, Supplementary Fig. S6**). Grids were prepared in the presence of ATPγS to prevent unwinding of the lagging strand. 2D class averages show clear density for the CUL1 stalk connected to FBH1, as well as diffuse bands of density emanating from FBH1 that correspond to the arms of the DNA fork (**Fig. 4a**). A 3.1-Å consensus 3D reconstruction shows the protein complex to consist of two regions—a “body” composed of FBH1, SKP1, and the N-terminal half of CUL1, and a more flexible “head” region composed of the C-terminal half of CUL1 and RBX1 (**Fig. 4b**). The local resolution of the body ranges from 2.8 Å at its core to 4.2 Å at its periphery, with a short stretch of ssDNA visible within the helicase domain (**Supplementary Fig. S6**). The resolution of the head region is 3.3 Å at the center of CUL1 and greater than 8 Å at the RBX1 end. Consistent with greater flexibility in the head region, 3D variability analysis showed bending of the CUL1 neck and multiple configurations of the CUL1/RBX1 head (**Supplementary Fig. S7**).

**Figure 4.**
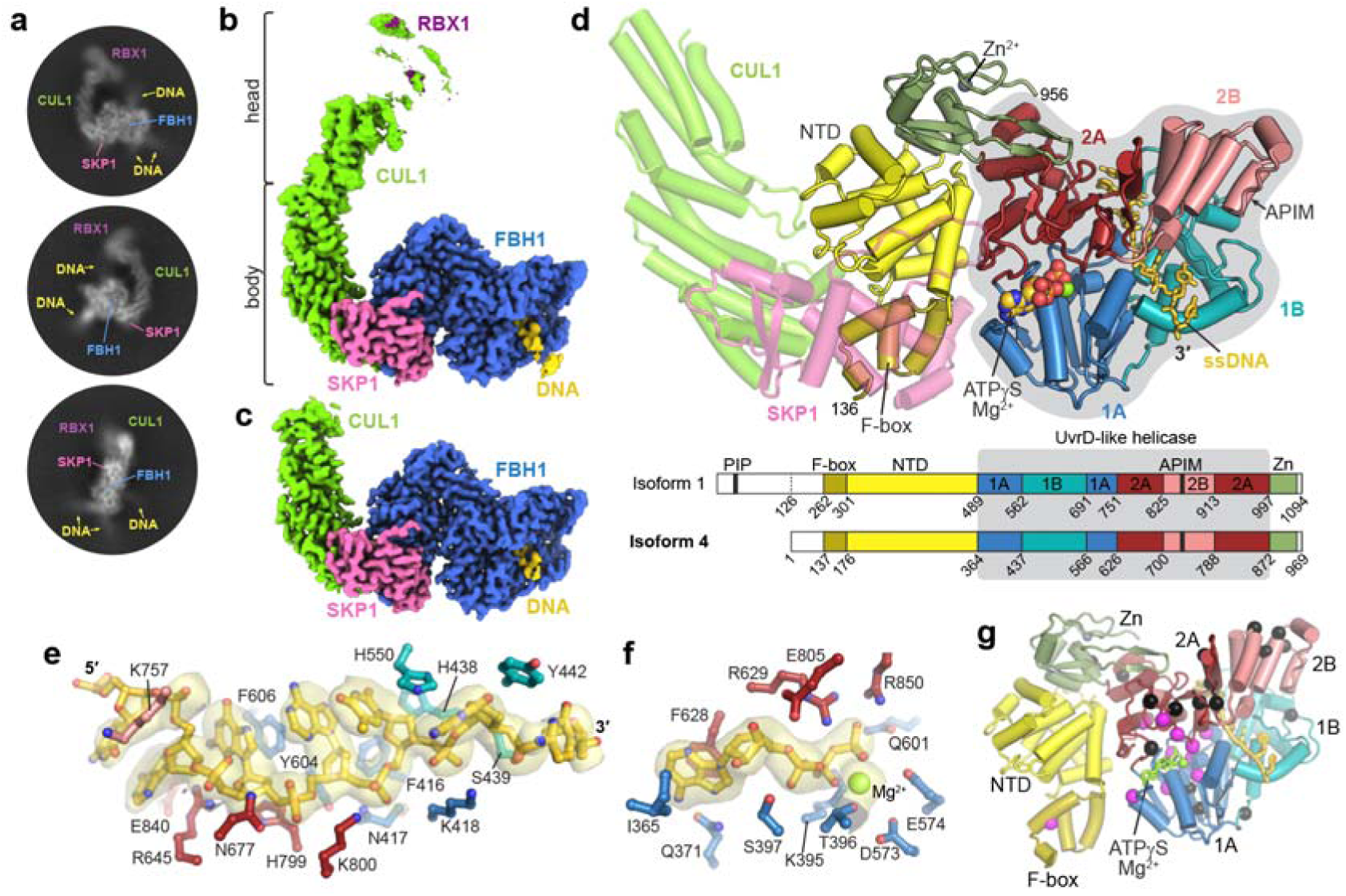
Cryo-EM structure of SCF^FBH1^. **a**. Representative 2D class averages. **b**. 3.1-Å consensus 3D reconstruction, colored according to subunit. **c**. 3.0-Å reconstruction of the body region. **d**. Atomic model of the body structure. CUL1, green; SKP1, magenta; FBH1, colored by subdomain, as specified in the primary structure schematic below. **e**. Details of the ssDNA in the helicase active site. Side chains are colored according to helicase subdomain as in panel d. Cryo-EM density is shown as a transparent surface. **f**. Cryo-EM density and side chain interactions of ATPγS/Mg^2+^. **g**. FBH1, colored by subdomain. Spheres indicate the Cα positions of FBH1 missense mutations associated with cancer (magenta) and other hereditary diseases (black), as identified by COSMIC^65^, HGMD^66^, and ClinVar^67^ databases.

To better characterize the details of FBH1, we performed focused 3D classification and refinement of the body region, which resulted in a 3.0-Å reconstruction (**Fig. 4c, Supplementary Fig. S5, Table S1**). The body structure contains residues 136-956 of FBH1 isoform 4, lacking the presumably unstructured 135 N-terminal residues immediately upstream of the F-box and 13 C-terminal residues. The structure contains the canonical interactions between the F-box and SKP1 observed in prior SCF structures^60, 61^. A unique α-helical N-terminal domain (NTD) immediately following the F-box and a C-terminal zinc ribbon further bridge the CUL1/SKP1 complex with the helicase domain (**Fig. 4d**). The helicase domain is oriented with the RecA-like catalytic 1A and 2A subdomains at the SKP1/NTD/zinc ribbon interface and the SF1-specific 1B and 2B subdomains facing away from the SCF complex. The map shows clear density for seven nucleotides of ssDNA and ATPγS/Mg^2+^ within the helicase domain (**Fig. 4e,f, Supplementary Fig. S5**). The sequence register and polarity of the DNA could be discerned from the density. As observed in other SF1 helicase structures^62, 63, 64^, the ssDNA makes backbone and base contacts with 1A, 2A, and 1B subdomains and runs 3′-5′ from 1A/1B to 2A/2B domains (**Fig 4d,e**), and the ATPγS nucleotide and catalytic Mg^2+^ ion make specific interactions with conserved helicase motifs at the 1A/2A interface (**Fig. 4f**). Highlighting the importance of the translocase activity of the protein, most disease-associated mutations^39, 65, 66, 67^ map to the ATP-binding interface and to the helicase 2B subdomain (**Fig. 4g**), which is an important regulatory element in SF1 helicases^20, 62, 63, 64, 68, 69, 70, 71, 72^.

### FBH1 contains a junction binding motif essential for fork reversal

Focused 3D classification and refinement to better characterize the weak DNA density evident in the 2D class averages resulted in a 10-Å “substrate” reconstruction of the FBH1 helicase and zinc ribbon domains bound to the fork (**Fig. 5a-c; Supplementary Fig. S5, Fig. S6**). Three cylinders of density representing parental, leading, and lagging arms branch off from the ends of the ssDNA that runs through the helicase domain (**Fig. 5c**). The lagging arm, representing the duplex to be unwound by the helicase, is connected to the ssDNA 5′-end. On the 3′-end, continuous density connects the ssDNA to the parental duplex, which itself is connected by continuous density to the leading arm. Major and minor grooves could be inferred from an alternating pattern of bulges and indentations in each cylinder of density (**Fig. 5c**). 3D variability analysis of the consensus reconstruction indicates multiple distinct motions of the DNA arms, in which the lagging arm pivots by ∼50° and the parental and leading arms, which form the fork junction, flex as a rigid body from their point of interaction with FBH1 (**Supplementary Fig. S7**). This limited range of motion at the junction places the leading arm of the fork in proximity to the APIM motif (733-KFIRR-737), which resides on a solvent exposed α-helix of the 2B subdomain such that it could interact with PCNA on the leading strand (**Fig. 5d**).

**Figure 5.**
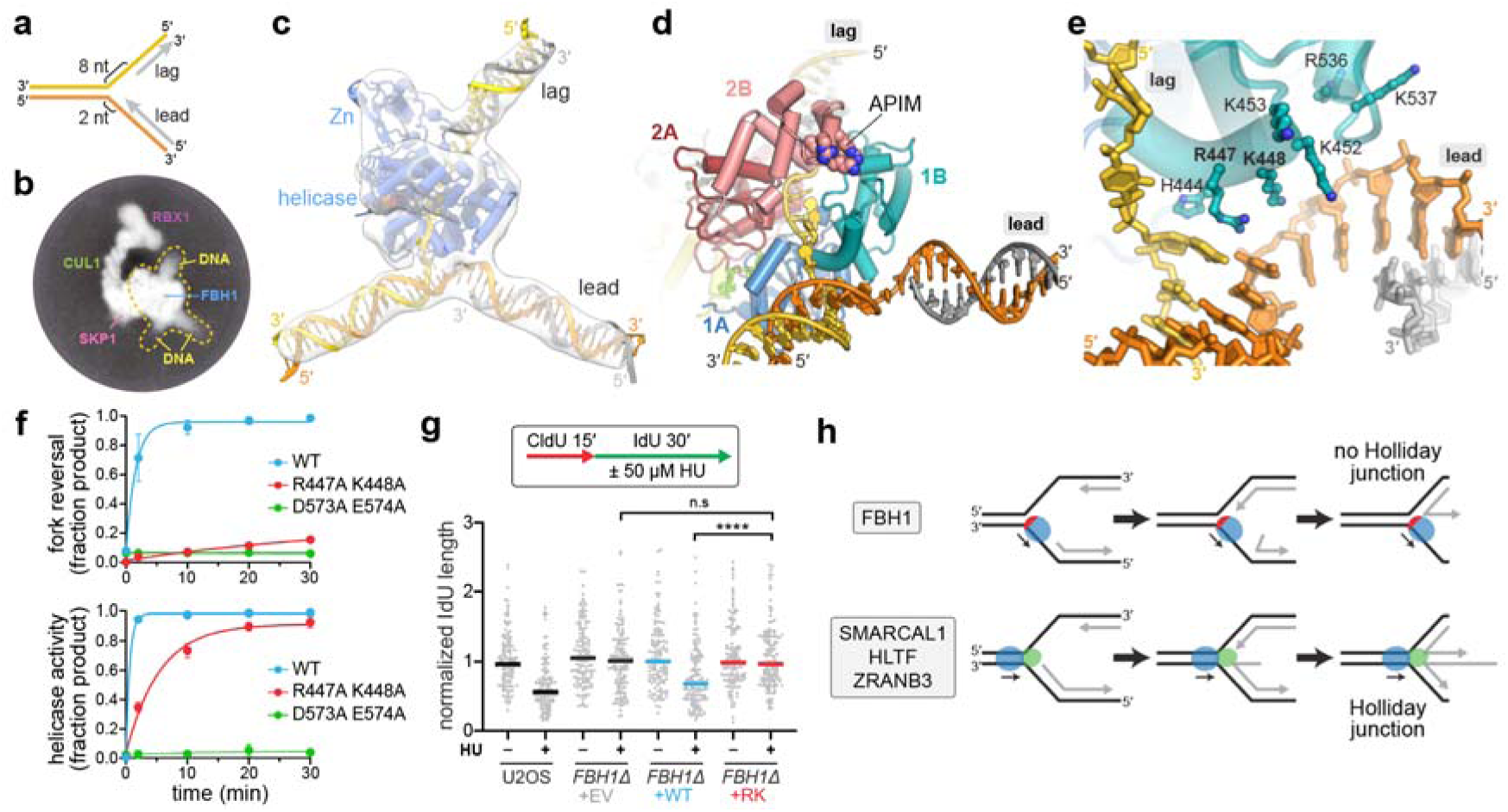
FBH1 contains a fork binding motif essential for fork reversal. **a**. Schematic of the DNA fork used in cryo-EM. **b**. Representative 2D class average showing diffuse density emanating from FBH1. The region defined by the 3D reconstruction in panel c is outlined. **c**. 10-Å substrate reconstruction. EM density is shown as a transparent white surface, protein is blue, template strands are orange (leading) and gold (lagging), and nascent strands are grey. **d**. Substrate structure colored according to helicase subdomain. APIM side chains are shown as spheres. **e**. Close-up of the junction binding motif of subdomain 1B. **f**. Fork reversal (top) and helicase (bottom) activities of SCF^FBH1^ wild-type and mutants. Representative gels are shown in Supplementary Fig. S8. **g**. DNA combing to measure replication for speeds in U2OS or U2OS *FBH1*Δ cells expressing the indicated FBH1 proteins. EV, empty vector; WT, wild-type; RK, R447A K448A. The experiment is representative of four biological repeats. **** *p* < 10^−3^ b a Kruskal-Wallis test; n.s., not significant. **h**. Model for fork reversal by FBH1 (top) and SNF2-family remodelers (bottom). The direction of the motor (blue) relative to the DNA is depicted by black arrows. Red, FBH1 junction binding motif; green, fork recognition domain of SNF2 remodelers.

The helicase 1B subdomain stabilizes the branch point of the junction through van der Waals contacts and a positively charged patch of residues that interact with both leading and lagging template strands (**Fig. 5e**). This junction binding motif is unique to FBH1; the 1B subdomains in other SF1 helicases lack the positive surface and contain extra structural elements that sterically clash with the fork junction in our structure. Within the junction binding motif, the guanidinium side chain of Arg-447 is stacked against the last base pair of the parental duplex, and the ε-amino group of Lys-448 interacts electrostatically with the phosphoribose backbone of the leading strand template. Substitution of Arg-447 and Lys-448 with alanines (R447A K448A, or “RK”) reduced fork reversal activity 25-fold relative to wild-type (**Fig. 5f, Supplementary Fig. S8, Table S2**), indicating the importance of the junction binding motif for fork reversal. This reduction of fork reversal activity is similar to that of the ATPase-dead Walker B mutant, although the RK mutation did not affect ATPase activity (**Supplementary Fig. S8**). In contrast to its effect on fork reversal, the RK mutant exhibited less than 10-fold reduction in helicase activity against a fork structure and no effect on unwinding a 3′-overhang substrate in our gel-based assay (**Fig. 5f, Supplementary Fig. S8, Table S2**). In the single-molecule assay, the RK mutant unwinding rates and dependence on applied force were comparable to wild-type, with maximum mean unwinding rate reaching 24 bp/s at 14 pN (**Supplementary Fig. S8**). The ssDNA translocation rates were also similar to wild-type, but the RK mutant exhibited a distinct force-dependent profile with faster rates at lower forces (peak rate of 42 nt/s at 6–8 pN that decreased to 33–35 nt/s at 10–14 pN). These data are consistent with the importance of the Arg-447 and Lys-448 interactions with the DNA junction, as loss of these interactions would lead to enzyme slippage and interfere with hairpin closure behind the enzyme. Thus, mutation of the junction binding motif severely impacts fork reversal, has a moderate effect on helicase activity, and does not interfere with ssDNA translocation by SCF^FBH1^.

To determine if the RK mutation affects the ability of FBH1 to perform fork reversal in cells, we utilized a single molecule DNA combing assay. Fork reversal in response to low doses of the replication stress agent hydroxyurea (HU) causes fork slowing^73, 74^, making measurement of fork speed a useful assay for reversal^7^. FBH1 knockout cells exhibited unrestrained fork speeds consistent with a fork reversal failure (**Fig. 5g, Supplementary Fig. S8**). Reintroduction of wild-type but not the RK mutant restored the fork slowing response to HU, consistent with the inability of the RK mutant to catalyze reversal (**Fig. 5g**).

These data indicate that the junction binding motif on subdomain 1B is essential for fork reversal *in vitro* and in cells, and support a model for fork reversal by FBH1 that is fundamentally different than that of SMARCAL, HLTF, and ZRANB3 (**Fig. 5h**). Because FBH1’s junction binding motif resides at the branch point of the two parental strands that reanneal during fork reversal, it stands to reason that FBH1 remains associated with the junction as it translocates on the lagging strand (as opposed to travelling away from the fork). A persistent interaction of FBH1 with the fork would facilitate duplex reannealing by threading the lagging strand template back into the junction, and may even destabilize the end of the leading duplex to facilitate its unwinding as the nascent strand would eventually contact the junction binding motif. Thus, whereas the dsDNA translocases SMARCAL1, HLTF, and ZRANB3 operate by pushing into the fork from the parental duplex side, FBH1 acts from behind the junction by pulling the lagging strand into the fork (**Fig. 5h**). Importantly, translocation on the lagging strand behind the fork would sterically prevent Holliday junction formation and branch migration, thus representing a distinct pathway—with different cellular outcomes—from SMARCAL1, HLTF, and ZRANB3^23^. This is consistent with our single-molecule data showing that SCF^FBH1^ cannot operate on the same fork substrates as the SNF2-family motors, and with the observation that the branch migration activity of the SNF2 motor protein RAD54L is essential for FBH1-mediated fork reversal in cells but dispensable for reversal by SMARCAL1 and HLTF^75^.

### FBH1 is a non-canonical SF1 helicase

The 2B subdomain is an important regulatory element in SF1 helicases that positions the DNA duplex to be unwound. Crystal structures of UvrD, PcrA, and Rep all show the 2B subdomain to exist in either an “open” conformation in the absence of DNA or a “closed” DNA-bound conformation, in which 2B scaffolds the DNA duplex to allow a strand-separation “pin” motif in subdomain 2A to access the ss/dsDNA junction (**Fig. 6a**)^62, 63, 64, 71, 76^. FBH1’s 2B subdomain is structurally divergent and does not contact the lagging duplex arm. Instead, the helical axis of the lagging arm is perpendicular to the FBH1 surface and ∼90° relative to the DNA duplexes in the SF1 structures (**Fig. 6b**). This conformation has major implications for FBH1’s ability to interact with RAD51, which we describe in the next section.

**Figure 6.**
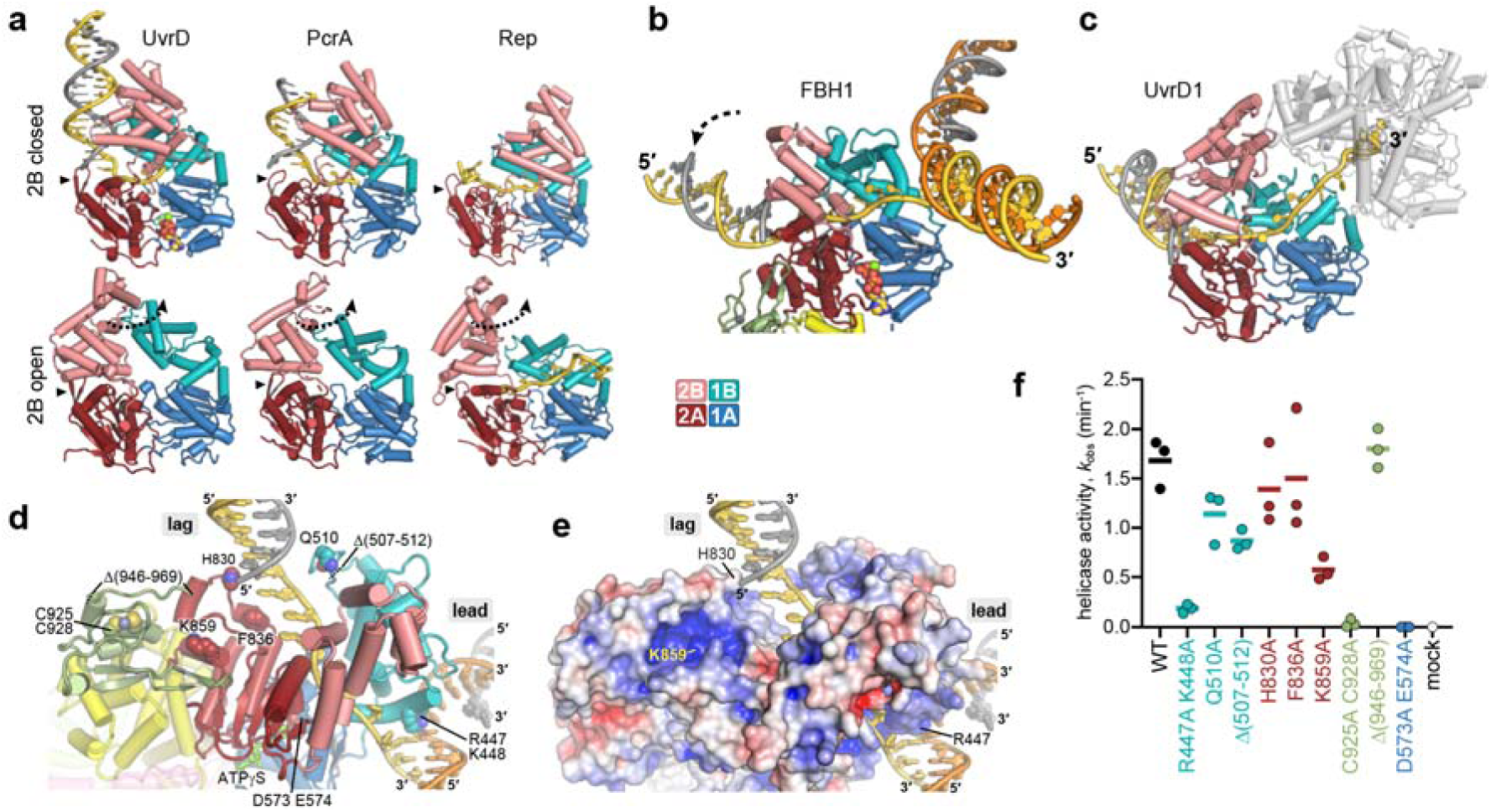
FBH1 is a non-canonical SF1 helicase. **a**. Crystal structures of SF1 helicases in DNA-bound (top) and unbound (bottom) states, colored by subdomain. Black triangles indicate the positions of the strand-separation pin motifs. The dotted arrows indicate the rotation of the 2B domain from open (unbound) and closed (bound) states. UvrD, PDB ID 2IS6 and 3LFU; PcrA, PDB ID 3PJR and 1PJR; Rep, PDB ID 1UAA. **b**,**c**. Structures of SCF^FBH1^ substrate complex (**b**) and *M. tuberculosis* UvrD1 dimer bound to DNA (PDB ID 9DES) (**c**) in the same orientation as structures in panel a. Polarity of the translocating strands (gold) is indicated. The leading UvrD1 subunit is colored by subdomain and the trailing subunit is grey. The dashed arrow in FBH1 highlights the rotated position of the lagging duplex relative to those in panel a. **d**. SCF^FBH1^ substrate complex, showing the location of FBH1 residues tested for helicase activity. FBH1 is colored according to subdomain. **e**. Same view as in panel d with protein depicted as an electrostatic potential surface (blue, positive; red, negative). **f**. Helicase activities of SCF^FBH1^ mutants, colored according to their location within the helicase. Rate constants are derived from single-exponential fits to kinetic data (see Supplementary Fig. S8). Bars indicate mean values (n=3). Mock, no-enzyme control.

Biochemical experiments have indicated that SF1 helicases are activated by interaction with another protein that repositions the 2B subdomain^68, 69, 70, 77^, suggesting that the closed conformation and position of the DNA duplex observed in the monomeric crystal structures may not represent an active state of the enzyme. Recently, a cryo-EM structure of a dimeric form of *M. tuberculosis* UvrD1, in which the two subunits are covalently linked via their 2B subdomains, showed an alternate, presumably active conformation that differs from either open or closed conformation observed in the crystal structures (**Fig. 6c**)^78^. The trajectory of the DNA duplex ahead of the leading (unwinding) UvrD1 subunit is nearly identical to that of the lagging arm in FBH1, suggesting that the SCF^FBH1^ structure represents an active state of the enzyme. Furthermore, the position of UvrD1’s 2B subdomain at the ss/dsDNA junction is influenced (i.e., UvrD1 is activated) by its interaction with the trailing subunit. Interestingly, the position of the fork junction in SCF^FBH1^ is in the same location as the trailing UvrD1 subunit (**Fig. 6c**), suggesting that FBH1 is activated by the fork through a physical relay across junction binding 1B and regulatory 2B subdomains.

FBH1 lacks the SF1 strand-separation pin, which implies that FBH1 employs an alternative mechanism of unwinding. The resolution of the substrate structure was not sufficient to discern the details between FBH1 and the 5′-end of the nascent lagging strand (**Fig. 5c**), preventing a clear model for strand displacement. We therefore tested the effects of protein motifs in proximity to the lagging arm for their effects on strand displacement from the fork substrate via mutagenesis (**Fig. 6d-f, Supplementary Fig. S8, Supplementary Table S2**). Deletion of residues 507-512, which form a loop on subdomain 1B that contacts the lagging arm, or alanine substitution of Gln-510 at its tip, reduced activity by 30-50% relative to wild-type, consistent with this motif stabilizing the lagging duplex during unwinding. On the other side of the DNA duplex, we surprisingly found little effect of mutating subdomain 2A residues His-830 and Phe-836, both of which are located at the ss/dsDNA junction. Similarly, deletion of the C-terminal 24 residues (Δ946-969) had no effect on helicase activity despite the proximity of the C-terminus to the lagging arm junction (although we could not resolve the C-terminal 13 residues in the structure).

The zinc ribbon domain just upstream from the C-terminus is unique to FBH1 and had a significant effect on helicase activity. The zinc ribbon interacts with helicase subdomain 2A though an extended β-hairpin (Fig. 4d, Fig. 6d). A double alanine mutant of two of the Zn-coordinating cysteine residues (C925A C928A) abrogated ATPase and helicase activities, suggesting that the mechanical motions of the helicase core are dependent on the zinc ribbon, although we cannot rule out the possibility that the mutant destabilized the global protein fold. Moreover, the β-hairpin forms one side of a deep, positively charged cavity that resides close to the lagging arm (**Fig. 6d,e**). Alanine substitution of Lys-859 at the back of this cavity reduced helicase activity 67% relative to wild-type, which was the second largest effect of any mutant we tested that still retained ATPase activity (**Fig. 6e, Supplementary Fig. S8**). It is unclear how the cavity impacts FBH1 activity, although it is tantalizing to speculate that it may capture the 5′-end of the displaced strand, as the size and charge of the cavity is appropriate for the triphosphate moiety present at the 5′-end of an Okazaki fragment. Taken together, the mutagenesis data indicate that residues on the leading edge of subdomains 1B and 2A work together with the zinc ribbon to facilitate strand separation. The relatively modest effect of these mutants on helicase activity suggests that translocation is more important than helicase activity for FBH1 function, consistent with the observation that strand displacement is not required for fork reversal by SCF^FBH1^ (**Fig. 3c**).

### A model for RAD51 filament disassembly

FBH1 acts as an anti-recombinase by displacing RAD51 from chromatin^27^ through both translocation-driven mechanical disassembly of RAD51 filaments^36, 37^ and ubiquitination of RAD51^33^. Our structure provides insight into how SCF^FBH1^ regulates DNA association of RAD51 (**Fig. 7**). Focused 3D classification and refinement of the head region of the SCF complex afforded a 3.9-Å reconstruction of the C-terminal half of CUL1 and the N-terminal region of RBX1 (**Fig. 7a**). However, the dynamic protein-protein interactions in this region did not allow for unambiguous placement of RBX1 other than the β-strand (residues 21-36) that is engulfed by CUL1. Nonetheless, combining the head structure with the body and substrate structures enabled construction of a nearly complete SCF^FBH1^ complex. This experimentally determined composite reconstruction revealed that the RBX1/CUL1 head region resides less than 45 Å from the lagging strand, which would place the ubiquitin ligase machinery in proximity of the lagging strand (**Fig. 7b**). Indeed, construction of a theoretical ubiquitin ligase-substrate model by docking RBX1, charged E2∼Ub, and the substrate of ubiquitination from the cryo-EM structure of CRL1^β-TRCP^/IκBα^61^ onto our SCF^FBH1^ structure places the ubiquitination machinery directly in the path of lagging strand (**Fig. 7c**). This model suggests that SCF^FBH1^ interacts with RAD51 bound to the lagging strand. This configuration would not be possible if the lagging strand were in the closed conformation observed in other SF1 structures. It is not clear whether SCF^FBH1^ ubiquitylates RAD51 as a filament or free in solution. Thus, SCF^FBH1^ may reduce chromatin-associated RAD51 by either (i) preventing reassociation of RAD51 with DNA via K64 ubiquitination^33, 79, 80^ after mechanical disassembly of the filament via FBH1 translocation or (ii) disassembling filaments through ubiquitination of subunits (**Fig. 7d**). Regardless, the major functional implication of our model is that RAD51 forms filaments on the lagging strand of a stalled fork during replication stress.

**Figure 7.**
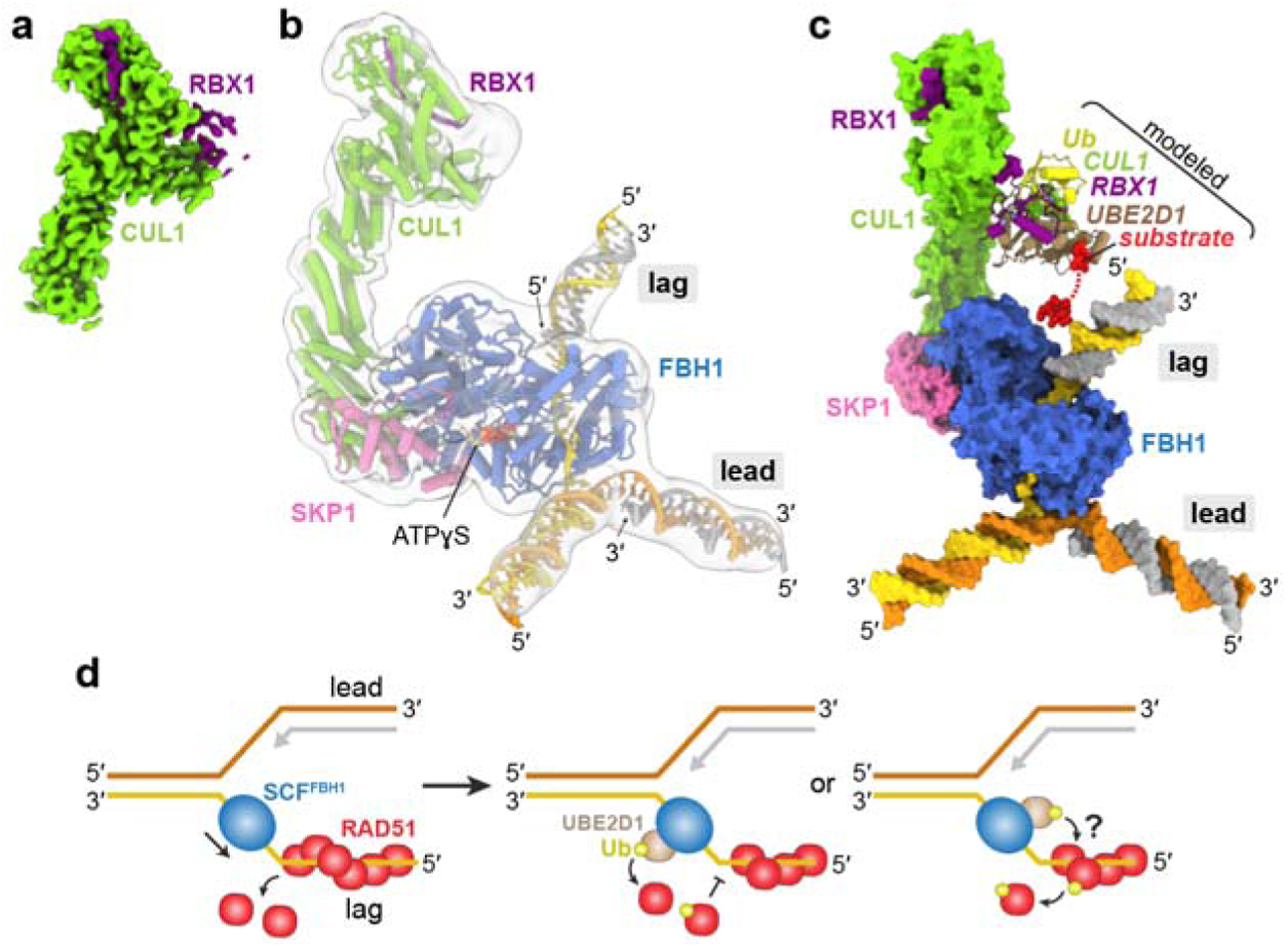
Model for RAD51 filament disassembly. **a**. 3.9-Å 3D EM reconstruction of the head region. **b**. Experimentally determined SCF^FBH1^/DNA structure (colored ribbons) docked into a lowpass-filtered composite EM map (transparent surface) created by combining the consensus, head, body, and for reconstructions. The molecular model is colored by protein subunit, parental DNA strands are orange (leading) and gold (lagging), and nascent strands are grey. **c**. Theoretical model of the SCF^FBH1^ ubiquitin ligase/substrate complex. The experimentally determined SCF^FBH1^/DNA structure is shown as a molecular surface. The E2∼Ub/substrate subcomplex composed of the RBX1 C-terminal domain, UBE2D1, ubiquitin (Ub), and Ub target substrate is modeled from PDB ID 6TTU and shown as ribbon and CPK spheres. **d**. Schematic for RAD51 filament disassembly by mechanical displacement (left), ubiquitination of free RAD51 (center), and ubiquitination of filament subunits (right).

While further studies will be needed to understand how FBH1, RAD51, and the other fork reversal translocases cooperate to promote reversal, one possibility is that FBH1 removes RAD51 when it is bound to the lagging template strand (**Fig. 7d**). In this scenario, RAD51 on the lagging strand would be an obstacle to reversal in the absence of FBH1. This hypothesis is consistent with the positioning of FBH1 and mirrors the need for RADX—another protein that destabilizes RAD51 filaments^81^—for fork reversal in the presence of persistent replication stress. Another hypothesis is that FBH1 acts on a RAD51-generated paranemic junction behind the CMG helicase, as previously proposed^82^. Finally, FBH1 could also initiate fork reversal in the absence of CMG since the presence of CMG would presumably preclude binding of FBH1 to the fork junction. These models are consistent with FBH1 initiation of reversal by unwinding the lagging strand (Fig. 5h). The genetic requirements of multiple translocases, the FBH1 helicase, RAD51, and other regulatory proteins for fork reversal indicate that much more work will be needed to fully understand the mechanism(s) of this complex replication stress tolerance pathway.

## Methods

### Protein expression and purification

Purified SCF^FBH1^ was produced using baculovirus infected insect cells using the Bac-to-Bac system (ThermoFisher Scientific). Genes for human FBH1 isoform 4 (UniProt Q2TAK1), SKP1, CUL1, and RBX1 were cloned into a p438-C insect cell expression vector^83^, which introduced a human rhinovirus (HRV) 3C protease cleavable N-terminal 6xHis-MBP tag to FBH1. Production of recombinant baculovirus and expression of SCF^FBH1^ genes was carried out in Sf9 cells. For expression, Sf9 cells were infected with amplified baculovirus at a MOI of 2 and harvested after 70 h. Insect cell pellets were resuspended in Ni buffer (20 mM HEPES pH 7.5, 500 mM NaCl, 10% glycerol, 20 mM imidazole, 1 mM TCEP) and lysed via dounce homogenization. The insoluble material was pelleted by centrifugation at 50,000 ×g for 30 min. Clarified lysate was applied to HisPur Ni-NTA resin (Thermo Fisher Scientific) and subsequently washed with additional Ni buffer to remove unbound contaminants. Bound proteins were eluted with Ni buffer containing 250 mM imidazole. HRV 3C protease was added to the elution to a final concentration of 25 ng/µL and the sample dialyzed against Heparin buffer (20 mM HEPES pH 7.5, 100 mM NaCl, 10% glycerol, 1 mM TCEP) overnight at 4°C. Following dialysis, the sample was centrifuged at 4,000 ×g for 10 min before loading onto a 5-mL Heparin HiTrap column (Cytiva) equilibrated in Heparin buffer. After washing, proteins were eluted over a 60-mL gradient from 0.1–1 M NaCl. Fractions containing the SCF^FBH1^ complex were pooled, concentrated to 10 mg/mL, and flash frozen in liquid nitrogen.

### DNA substrate preparation

All oligodeoxynucleotides used in this study are listed in Table S3, and strand combinations annealed to form specific substrates are described in Table S4. Oligos were annealed from 98° to 25°C over 60 min in 1× SSC buffer (15 mM sodium citrate pH 7.0 and 150 mM NaCl). 6-carboxyfluorescein (FAM)-labeled substrates were used in DNA binding assays. The duplex containing a 3′-overhang was generated by annealing *FAM40* to *FAM40_3*′*oh_8* in a 1:1.2 molar ratio. Heterologous forks were prepared by first annealing the leading (*FAM40*/*FAM40_lead_2gap* or *FAM40*/*FAM40_lead_8gap*) and lagging (*F20.40*/*FAM40_lag2gap* or *F20.40*/*FAM40_lag8gap*) arms separately, followed by annealing the duplexed arms at 37°C using a 1.2-fold molar excess of unlabeled duplex. Helicase and fork reversal assays contained ^32^P-labeled substrates. Oligos were 5′-labeled by incubating with ^32^P-γATP and polynucleotide kinase (NEB) at 37°C for 30 min, followed by heat inactivation at 65°C for 10 min and purification via a G-25 column (GE Healthcare). The 3′-overhang substrate used in the helicase assay was generated by annealing ^32^P-labeled *9oh-helicase* to *53_5gap* using a 1.2-fold molar excess of unlabeled oligonucleotide. Heterologous forks utilized in the helicase assay were generated by annealing ^32^P-labeled leading arm (*54/56*) to a two-fold molar excess of the lagging arm (*52/53_8gap*). Homologous forks used in the fork reversal assay were generated by annealing the ^32^P-labeled leading (*48/50*, *48/50_5gap*, or *A/B*) and lagging (*52/53*, *52/53_5gap*, or *D*) halves of the fork separately, followed by incubation of unlabeled and labeled duplexes in a 2:1 molar ratio at 37°C for 30 min. Fork reversal substrates were PAGE purified using a 6% 0.5X TBE DNA Retardation Gel (Invitrogen). Bands were excised from the gel and electroeluted in 0.5× TBE, concentrated using a 10K Amicon-Ultra 0.5 ml centrifugal filter, and stored at −20°C. The heterologous fork used for the cryo-EM structure was generated by annealing the four EM oligos shown in Table S1.

### DNA binding

DNA binding affinity was determined using an electrophoretic mobility shift assay in which varying concentrations of SCF^FBH1^ (0–500 nM) was incubated with 100 nM 5′-FAM labeled substrate (Table S3 and S4) in binding buffer (20 mM HEPES pH 7.5, 100 mM NaCl, 0.5 mM TCEP, and 0.02% NP-40). Reactions were incubated at 4°C for 30 min electrophoresed at 200V for 1 h in a 5% 79:1 (acrylamide:bis-acrylamide) 1× TBE gel containing 5% (v/v) glycerol. Gels were visualized using a ChemiDoc MP (Bio-Rad) and band intensities were quantified using GelAnalyzer (www.gelanalyzer.com). Fraction bound was defined as *I*_bound_ / (*I*_bound_ + *I*_free_), where *I*_bound_ and *I*_free_ are the intensities of bands corresponding to SCF^FBH1^•DNA and free DNA, respectively. Statistical analysis was performed using Prism10 (GraphPad).

### DNA helicase assay

Helicase reactions were carried out in solution at 21°C for 30 min and contained 5 nM SCF^FBH1^, 1 nM ^32^P-labeled fork substrate (Table S3 and S4), and 10 nm trap oligonucleotide (Table S3) in reaction buffer (20 mM Tris pH 8.0, 50 mM NaCl, 5 mM MgCl_2_, 1 mM TCEP, 2 mM ATP, and 100 ug/ml BSA). Reactions were stopped by the addition of Proteinase K (Sigma) and electrophoresed on an 8% 19:1 (acrylamide:bis-acrylamide) 1× TBE gel at 10 W for 1.5 h. Gels were phosphorimaged on a Typhoon RGB imager (Cytiva), band intensities were quantified using ImageQuantTL (Cytiva), and analyzed using Prism10 (GraphPad).

### Fork reversal assay

Fork reversal reactions were carried out as previously described^13^, with minor modifications. Briefly, reactions were performed at 21°C or 37°C and contained 5 nM SCF^FBH1^, 1 nM ^32^P-labeled fork substrate (Table S3 and S4), 2 mM ATP, and 100 ug/ml BSA in either 40 mM Tris pH 8.0, 50 mM NaCl, 5 mM MgCl_2_, and 1 mM TCEP (SCF^FBH1^) or 20 mM Tris pH 7.76, 10 mM KCl, 2 mM MgCl_2_, and 1 mM DTT (HLTF). Reactions were terminated by the addition of Proteinase K (Sigma) and electrophoresed on an 8% 19:1 (acrylamide:bis-acrylamide) 1× TBE gel at 10 W for 1.5 h. Gels were phosphorimaged and quantified as described above.

### Magnetic tweezers assays

#### DNA substrates

The hairpin substrate used in the helicase/translocase and annealing assays was constructed based on a previous design ^84^ with some modifications. The substrate consists of a 74-bp dsDNA fragment ligated to a 126-bp highly biotinylated dsDNA handle, a 1,238-bp hairpin with a dT_4_ loop, and a 146-bp 3′-digoxigenin labeled dsDNA tail (Table S5). The digoxigenin labeled dsDNA fragment is connected to the hairpin through a ssDNA dC_30_ region. The fork substrate used in the reversal/branch migration assay was based on a previously published design ^54, 59^ but fabricated differently. The substrate consists of two identical dsDNA fragments arranged in an inverted orientation and connected by a short hairpin, forming a three-way junction capable of branch migration. The spontaneous branch migration is avoided by introducing a mismatch of 1 bp at the beginning of the hairpin. The structure is ligated to two highly labeled DNA fragments, one with digoxigenins and the other with biotins, of 997 bp and 152 bp, respectively, used as immobilization handles. Details for construction of the substrates are described in the Supplementary Material.

#### Magnetic tweezers instrument

The magnetic tweezers setup used in this work was similar to previously described systems^85, 86^, with variations introduced depending on the specific experiment—helicase/translocase versus fork reversal/branch migration. The core setup consisted of a pair of vertically aligned permanent NdFeB magnets positioned above a flow chamber mounted on an inverted microscope. DNA molecules tethered on one end to 1-μm diameter MyOne C1 superparamagnetic beads were introduced into the chamber and immobilized on its lower surface on the other end. The magnets enabled force application by attracting the superparamagnetic beads, thereby stretching the tethered DNA molecules. Bead position was tracked at 120 Hz using a CCD camera coupled to a high-magnification oil-immersion objective. The applied force, which depended on the distance between the magnets and the sample, was estimated from the Brownian fluctuations of the beads ^85^. DNA extensions were measured by analyzing images captured at different focal planes.

For fork reversal experiments, which required only moderate forces, the magnets (Supermagnete) were separated by a 1-mm gap. Flow chambers were assembled by sandwiching a single layer of parafilm cut to define the flow channel between two glass coverslips. The upper coverslip was perforated with two small holes to allow fluid exchange (inlet and outlet). This configuration allowed the application of forces up to ∼7 pN to 1 μm beads. For helicase/translocase experiments, which required higher forces, the setup was adapted to achieve stronger magnetic field gradients. Two small, vertically aligned N52 magnets (HKCM, 9964-5112) with a 0.3-mm inter-magnet gap were employed. To minimize the distance between the beads and the magnets, chambers were constructed using double-sided adhesive tape (Adhesive Research, 92712) to define the flow channel. The upper coverslip was replaced with a thin plastic Mylar film. This configuration enabled the application of forces up to ∼25 pN to 1 μm MyOne C1 beads.

#### DNA Immobilization

To immobilize DNA molecules, the flow chambers were first functionalized by passive adsorption of anti-digoxigenin antibodies onto the polystyrene-coated lower coverslip. Specifically, chambers were incubated overnight at 4 °C with 100 ng/μl anti-digoxigenin antibody (Bio-Rad). The polystyrene coating was applied to the lower coverslips during chamber fabrication. Before introducing DNA-bead complexes, the antibody-coated chambers were passivated with 1 mg/ml bovine serum albumin (New England Biolabs) in PBS buffer (PBS-BSA) for 45 min at room temperature to minimize nonspecific binding.

To prepare the DNA-bead complexes, 0.3–0.5 ng of DNA in TE buffer (10 mM Tris-HCl, pH 8.0, 1 mM EDTA) was mixed with 5 μl of 1 μm-diameter streptavidin-coated magnetic beads at stock concentration. Prior to use, the beads were washed in PBS-BSA. The mixture was incubated for 10 min at room temperature on a rotator to allow binding of the biotinylated DNA handles to the streptavidin beads. Following this, 25 μl of PBS-BSA was added and the mixture was incubated for an additional 20 min. After incubation, 5–10 μl of the DNA-bead mixture was diluted in PBS-BSA and introduced into the passivated flow chamber. This allowed the digoxigenin-labeled end of the DNA to bind to the surface-immobilized anti-digoxigenin antibodies. After an incubation of 8 min, unbound beads were removed by flushing the chamber with PBS.

#### Experimental

Helicase and translocation activity were measured using the DNA hairpin substrate. SCF^FBH1^ (100 pM) was introduced into the flow chamber in FBH1 Buffer (20 mM Tris-HCl (pH 8), 50 mM NaCl, 2 mM ATP, 5 mM MgCl_2_, 1 mM DTT). The applied force was maintained below the hairpin’s mechanical opening threshold (15 pN), and DNA extension was monitored throughout. Rewinding activity assays were also conducted with the DNA hairpin substrate, using either 40 nM SCF^FBH1^ in FBH1 Buffer or 10 nM RecG in RecG Buffer (25 mM TrisOAc (pH 7.5), 150 mM KOAc, 1 mM ATP, 1 mM MgCl_2_, 1 mM DTT). The applied force was maintained at approximately 15 pN, corresponding to the mechanical opening threshold of the hairpin. Under these conditions, extension oscillations consistent with spontaneous hairpin opening and closing were observed, suggesting the presence of a short duplex region near the hairpin loop in the substrate. DNA extension was then continuously monitored. Reversal assays were performed using the fork substrate, with DNA extension monitored continuously for 90 min, using 40 nM SCF^FBH1^ in FBH1 Buffer, 40 nM HLTF in HLTF Buffer 2 (40 mM Tris-HCl (pH 7.5), 50 mM NaCl, 1 mM ATP, 2 mM MgCl_2_, 2 mM DTT), or 10 nM RecG in RecG Buffer.

#### Data analysis

Unwinding and translocation rates were computed from the DNA hairpin assay data using custom Python 3 scripts (https://github.com/Moreno-HerreroLab/MT_Helicase_HairpinData_Analysis). First, the extensions were transformed to unwound bp using ssDNA force extension curves acquired in either FBH1 or RecG Buffer and using a blocking oligo^87, 88^ (**Supplementary Fig. S2**). Time courses were filtered by applying the Chung-Kennedy non-linear filter^89^ with parameters K=100, M=5, and p=1 two times. The protein almost never paused during unwinding or translocation. Thus, pauses were eliminated from the analysis so that the unwinding rates obtained were not perturbed by them. Then, the instantaneous rate at each time-point of every unwinding or translocation event was obtained and the mean of the derivatives of each event was taken as a point for the rate-force dependency analysis. The graphs and statistical analysis were performed using OriginPro software.

### Cryo-EM

#### Sample preparation and data processing

SCF^FBH1^ (20 µM) was incubated with the EM fork substrate in a 1:1.2 protein:DNA molar ratio for 30 min on ice. The complex was then incubated with BS^3^ crosslinker (ThermoFisher) at a final concentration of 0.5 mM on ice for an additional 30 min, after which the reaction was quenched by the addition of Tris pH 8 to a final concentration of 50 mM. The crosslinked sample was injected onto a Superdex 200 10/300 column (Cytiva) equilibrated in SEC buffer (20 mM HEPES pH 7.5, 150 mM KCl, 5 mM MgCl_2_, 1 mM TCEP) to remove any aggregated protein or unbound DNA. Fractions containing SCF^FBH1^ were pooled and concentrated to 1 mg/mL. Prior to freezing grids, SCF^FBH1^ was diluted to 0.6 mg/mL and ATPγS was added to a final concentration of 2 mM. Grids were prepared by applying 2.5 µL of SCF^FBH1^ to glow discharged UltrAuFoil R1.2/1.3 Au300 grids (Quantifoil) followed by blotting and plunge freezing in liquid ethane using a Vitrobot Mark IV (Thermo). Videos of dose-fractionated frames were collected from two grids in two separate imaging sessions on a Titan Krios electron microscope (Thermo) operating at 300 kV with an overall dose rate of 51.8 or 54.5 e/Å^2^ and a magnification of ×105,000 using a Gatan K3 direct electron detector operating with a Gatan imaging filter.

Preprocessing of the raw videos, consisting of patch motion correction and patch CTF estimation, as well as all initial data-processing steps, were performed in cryoSPARC^90^. Micrographs with a CTF resolution of worse than 6 Å were discarded. Particles were picked from the remaining 15,611 micrographs using Topaz^91^ and a model trained from a small set of manually picked particles. Particles were then extracted in a downsampled 100-px box (3.272 Å/px) and sorted by 2D classification. Classes exhibiting clear secondary structural elements and/or rare views, consisting of 881,920 of 2,286,823 particles, were selected. Using the downsampled particles, a preliminary 3D reconstruction was generated through ab initio reconstruction and non-uniform refinement. Particles were then recentered and re-extracted without downsampling in a 540-px box (0.818 Å/px), and non-uniform refinement was repeated using the 829,277 re-extracted particles. All subsequent data-processing steps were performed in RELION^92^. Focused 3D refinement with Blush regularization^93^ was performed to improve the consensus reconstruction. Regions with weak density, including much of CUL1/RBX1 and the entirety of three DNA duplex arms of the fork, were further improved by focused 3D classification without alignment, using masks for the head, the body, and the substrate. A single class was chosen for each region, consisting of 156,479, 181,442, and 53,451 particles, respectively. For the substrate region, density for the DNA duplexes was further improved by particle subtraction and a second round of 3D classification without alignment, from which a single class consisting of 7,718 particles was selected. These subtracted particles were exchanged with the corresponding original (presubtraction) particles for further processing. For all three regions, final reconstructions were generated using focused 3D refinement with Blush regularization. Final maps were sharpened using DeepEMhancer^94^.

#### Model building and refinement

An initial model of FBH1 was generated using AlphaFold3^95^. Initial models of SKP1, CUL1, and RBX1 were obtained from PDB accession 1LDK^60^. All models were fit as rigid bodies into corresponding density maps using ChimeraX^96^. Models were further modified in Coot^97^ and were subjected to restrained atomic refinement against unmodified/unsharpened maps in PHENIX^98^. FBH1 (residues 136-956), SKP1 (2-161), and CUL1 (16-300) were first refined against the body map, and CUL1 (301-776) and RBX1 (21-36) were first refined against the head map. FBH1 (364-956) from the refined model, along with ideal B-form DNA duplexes generated in Coot, were then fit as rigid bodies into the substrate map. Because of the low resolution of the reconstruction, FBH1 was refined using reference restraints from the starting model and the DNA duplexes were refined using base-stacking and base-pairing restraints. Neither the first two nucleotides of the lagging strand nor the last two nucleotides of the leading strand could be confidently modeled. To generate a structure of the complete complex, the three partial models were docked into the consensus map and then combined into a single structure. The combined structure was refined using reference restraints from each of the partial structures. All refined models were validated with MolProbity^95^.

### DNA combing assay

U2OS cells were cultured in Dulbecco’s modified Eagle’s medium (DMEM) with 7.5% fetal bovine serum. *FBH1*Δ cells were described previously^23^. FBH1 expression vectors with isoform 4 of the FBH1 cDNA were generated with pLNCX2 with an 3XHA tag (pWL54, wild-type; pWL83, R572A,K573A). Cells were plated 24 hr before treatment to reach 50-70% confluence. To label newly synthesized DNA, cells were first treated with 5-chloro-2′-deoxyuridine (CldU, 25 μM, 15 minutes) (Sigma-Aldrich, C6891) and then 5-iodo-2′-deoxyuridine (IdU, 120 μM, 30 minutes) (Sigma-Aldrich, l7125). Cells were treated as indicated with hydroxyurea (Millipore Sigma, H8627). Approximately 500,000 cells were embedded in 1.5% low-melting agarose plugs in phosphate-buffered saline (PBS) and digested overnight in 0.1% sarkosyl, proteinase K (2 mg/ml), and 50 mM EDTA (pH 8.0) at 50°C. Plugs were washed in TE (10mM Tris pH=8.0, 1mM EDTA), transferred to 100 mM MES (pH 5.7), melted at 68°C, and digested with 1.5 U of β-Agarase I (New England Biolabs, M0392S) overnight at 42°C. DNA was combed on silanized coverslips using a GenomicVision combing apparatus. The DNA was stained with antibodies recognizing IdU (BD, 347580) and CldU (Abcam, ab6326) for 1 hr, washed in PBS, and probed with fluorescent secondary antibodies (Alexa Fluor 488, A28175, Alexa Fluor 555, A-21434) for 30 min. Images were obtained using a Nikon Ti2 microscope. Fiber lengths were measured manually from blinded samples.

## Data availability

Structures were deposited in the Protein Data Bank under accession codes 9XZJ (SCF^FBH1^/DNA), 9XZK (CUL1_301-776_/RBX1), 9XZL (FBH1/SKP1/CUL1_16-300_/ssDNA), and 9XZM (FBH1_364-956_/DNA). Maps were deposited in the Electron Microscopy Data Bank under accession codes EMD-72358 (consensus), EMD-72359 (head), EMD-72361 (body), and EMD-72632 (substrate). Source Data are provided with this paper.

## Acknowledgments

We thank the staff of the Vanderbilt Cryo-EM facility for assistance. This work was funded by the National Institutes of Health (R35GM136401 to B.F.E., R01GM116616 to D.C., and P01CA092584 to B.F.E. and D.C.). S.K.H. was supported by the Cellular, Biochemical, and Molecular Sciences Training Program (NIH T32GM137793). Work in the FMH laboratory was supported by grants PID2023-146255NB-I00, funded by MICIU/AEI/10.13039/501100011033 and FEDER, EU, and TEC-2024/TEC-158, funded by the Autonomous Region of Madrid and the European Social Fund and the European Regional Development Fund. JMG is supported from the Spanish Ministry of Universities through an FPU grant (FPU21/03892). The Vanderbilt Cryo-EM facility was funded in part by NIH S10OD030292. The content is solely the responsibility of the authors and does not necessarily represent the official views of the National Institutes of Health.

## Author contributions

Conceptualization: CJS and BFE; Data curation: EAM and BFE; Formal analysis: BHG, JMG, EAM, EMP, SKH, CJS, DC, FMH, and BFE; Investigation: BHG, JMG, EAM, SKH, and CJS; Project administration and Supervision: BFE and FMH; Resources: CAR and MST; Visualization: BHG, JMG, EAM, EMP, SKH, DC, FMH, BFE; Validation and Writing: all authors.

