## Supplementary for "Structural basis for fork reversal and RAD51 regulation by the SCF ubiquitin ligase complex of F-box helicase 1"

### Supplementary Methods

#### *Construction of magnetic tweezers substrates*

Oligodeoxynucleotide sequences for preparing substrates are described in Table S5.

DNA hairpin. The dsDNA hairpin used in helicase/translocase experiments was obtained by PCR amplification with Phusion High-Fidelity DNA Polymerase (Thermo Scientific) using Lambda DNA (NEB) as template and oligonucleotides that include different BsaI restriction sites on each side of the PCR fragment followed by purification (QIAGEN) as previously described[1]. This restriction site was selected to avoid nonspecific ligation products in the later steps. After BsaI digestion and purification, we obtained a dsDNA fragment of 1,204 bp with a homogeneous GC content and unique non-palindromic 5′-overhangs. A fork structure was formed by two partially complementary oligonucleotides, the *359.5-Phosphate BsaI* and the *293.Template_hairpin_30dC*. The two oligonucleotides were annealed by heating at 95°C for 5 min and cooling down to 20°C at a 1°C/min rate in hybridization buffer (10 mM Tris-HCl pH 8.0, 1 mM EDTA, 200 mM NaCl, 5 mM $MgCl_2$). The final fork structure contained a 5′-cohesive end compatible with one of the ends of the PCR fragment. The oligonucleotide *250.Loop hairpin* was self-annealed to create a short dsDNA hairpin with a cohesive end compatible with the other end of the PCR fragment. The fork structure and the short hairpin oligo (5-fold excess of each) were ligated overnight to either end of the 1,204-bp PCR fragment using T4 DNA ligase (NEB). The ligated DNA structure was annealed with 10-fold excess *360.3_BsaI_anneal_359* oligonucleotide (which is complementary to the *359.5-Phosphate BsaI* oligonucleotide), by heating at 95°C for 5 min and cooling down to 20°C at a 1°C/min rate in annealing buffer (10 mM Tris-HCl pH 7.5, 1 mM $MgCl_2$). After hybridization, a new BsaI 5′-cohesive end that was compatible with the short biotinylated dsDNA handle was created. A 2-nt gap remained next to the hairpin structure. The dsDNA handle was prepared by PCR using 200 µM final concentration of each dNTP (dGTP, dCTP, dATP), 140 µM dTTP and 66 µM Bio-16-dUTP (Roche) using the plasmid pSP73-JY0 as a template[2] followed by digestion with the restriction enzyme BsaI. Then, the DNA structure already hybridized with the *360.3_BsaI_anneal_359* oligonucleotide was ligated overnight with 10-fold excess of the biotinylated dsDNA handle. The ligated DNA construct was gel extracted to remove the excess oligonucleotides and purified (QIAGEN). Finally, the purified structure was annealed with 5-fold excess of the *253.Primer_for_Dig* oligonucleotide that was partially complementary to the oligonucleotide *293.Template_hairpin_30dC* by heating at 95°C for 1 min and cooling down from 80°C to 10°C at a 0.05°C/sec rate in annealing buffer. The digoxigenin labeling required to attach the DNA hairpin construct to a glass surface via anti-digoxigenin antibodies was then incorporated by filling in the overhangs with exonuclease-deficient DNA polymerase I Klenow Fragment (NEB) in the presence of dATP, dCTP and dUTP-digoxigenin (Roche) for 1 h at 37°C, followed by heat inactivation. A $dC_{30}$ ssDNA region remained as the primer was not completely extended due to the absence of dGTP. The completed DNA hairpin construct was ready to use in magnetic tweezer experiments without further purification. EDTA pH 8.0 was added to a final concentration of 1 mM. DNAs were never exposed to intercalating dyes or UV radiation during their production and were stored at 4°C.

Fork substrate. The design of a DNA substrate that mimics a stalled replication fork was based on the construct employed by Manosas et al[3, 4] but following a different fabrication method. While their construct was generated *in situ* in the reaction chamber by employing a T4 holoenzyme complex, our substrates were prepared beforehand. Two identical dsDNA fragments of 2 kbp arranged in an inverted orientation and connected by a short hairpin of 30 bp with a 4-nt loop

($dT_4$) were used to form a three-way junction capable of branch migration. A mismatch of 1 bp was included at the beginning of the hairpin to avoid spontaneous branch migration. The two dsDNA fragments of 2 kbp were obtained by PCR amplification with Phusion High-Fidelity DNA Polymerase (Thermo Scientific) using the plasmid pSP73-JY0 [2] as template and oligonucleotides that include KpnI and PspOMI restriction sites in one side of the PCR fragment and a BbvCI site in the other side. After purification (QIAGEN), the dsDNA fragment that would be connected to the digoxigenin-labeled handle was digested with Nt.BbvCI restriction enzyme (NEB) creating a 5′-overhang of 18 nt in one end. The dsDNA fragment that would be connected to the biotinylated handle was digested with Nb.BbvCI variant (NEB), creating a 3′-overhang of 15 nt on one end. This strategy to generate both 3′ or 5′-overhangs using nicking enzymes has been previously described[5, 6]. The 5′-overhang of 18 nt was annealed with 25-fold excess of oligonucleotide corresponding to the A-DIG side adaptor and the 3′-overhang of 15 nt was annealed with the A-BIO side adaptor by heating 10 min at 72°C and slowly cooling down to 42°C at a -0.1°C 25 $s^{-1}$ rate in annealing buffer followed by overnight ligation with T4 DNA Ligase (NEB). The complete 2 kbp-dsDNA branches were gel extracted and purified (QIAGEN). Then, the dsDNA fragment that would be connected to the digoxigenin-labeled handle was digested with PspOMI (NEB) and the dsDNA fragment that would be connected to the biotinylated handle was digested with KpnI (NEB), generating the overhangs to later ligate with the dsDNA handles. After digestion, the two branches were purified and mixed in equimolar ratio to subsequently perform the annealing between them through the DIG side adaptor and the BIO side adaptor, by heating as described above in the same annealing buffer. The oligonucleotide *250.Loop hairpin* was self-annealed by heating at 95°C for 5 min and cooling down to 20°C at a -1°C $min^{-1}$ rate in hybridization buffer to create a short dsDNA hairpin with a cohesive end compatible with the cohesive end formed once the DIG side adaptor and the BIO side adaptor were annealed between them. The dsDNA handles labelled with digoxigenins (997 bp) or with biotins (152 bp) were prepared by PCR (Table SX) including 200 µM final concentration of each dNTP (dGTP, dCTP, dATP), 140 µM dTTP and 66 µM Dig-11 dUTP or Bio-16-dUTP (Roche) using the plasmid pSP73-JY0 as template [2] followed by digestion with the restriction enzyme PspOMI or KpnI, respectively. In a final step, the two annealed branches were ligated overnight with the labelled handles together with a 10-fold excess of the self-annealed short dsDNA hairpin. The sample was ready for use without further purification. DNAs were never exposed to intercalating dyes or UV radiation during their production and were stored at 4˚C with 1 mM EDTA pH 8.0.

**Supplementary Table S1.** Cryo-EM data collection, processing, refinement, and validation statistics.

| Map name | **Consensus** | **Head** | **Body** | **Substrate** |
|---|---|---|---|---|
| Composition | $SCF^{FBH1}$/DNA | $CUL1^{301\text{-}776}$/RBX1 | FBH1/SKP1/CUL1 $^{16\text{-}300}$/ssDNA | $FBH1^{364\text{-}956}$/DNA |
| EMDB accession number | EMD-72358 | EMD-72359 | EMD-72361 | EMD-72362 |
| PDB accession number | 9XZJ | 9XZK | 9XZL | 9XZM |
| **Data collection and processing** | | | | |
| Magnification | ×105,000 | ×105,000 | ×105,000 | ×105,000 |
| Voltage (kV) | 300 | 300 | 300 | 300 |
| Electron exposure (e−/Å$^2$) | 51.8, 54.5 | 51.8, 54.5 | 51.8, 54.5 | 51.8, 54.5 |
| Defocus range (μm) | −2.0 to −1.0 | −2.0 to −1.0 | −2.0 to −1.0 | −2.0 to −1.0 |
| Pixel size (Å) | 0.818 | 0.818 | 0.818 | 0.818 |
| Symmetry imposed | C1 | C1 | C1 | C1 |
| Initial particle images (no.) | 2,286,823 | 2,286,823 | 2,286,823 | 2,286,823 |
| Particle images after 2D classification (no.) | 881,920 | 881,920 | 881,920 | 881,920 |
| Final particle images (no.) | 829,277 | 156,479 | 181,442 | 7,718 |
| Map resolution (Å) | 3.13 | 3.91 | 3.00 | 10.27 |
| FSC threshold | 0.143 | 0.143 | 0.143 | 0.143 |
| Map resolution range (Å) | 2.8–8.0 | 3.6–7.2 | 2.8–4.4 | >8.0 |
| **Structure refinement and validation** | | | | |
| Initial model used (PDB code) | 9XZK, 9XZL, 9XZM | 1LDK, AlphaFold3 | 1LDK, AlphaFold3 | 9XZL |
| Model resolution (Å) | 3.07 | 4.05 | 2.93 | 16.69 |
| FSC threshold | 0.5 | 0.5 | 0.5 | 0.5 |
| Map sharpening *B*-factor (Å$^2$) | −82.9 | −86.1 | −50.7 | −200 |
| Model composition (no.) | | | | |
| Non-hydrogen atoms | 16,522 | 3,985 | 10,111 | 7,341 |
| Protein residues | 1,723 | 492 | 1,231 | 593 |
| DNA nucleotides | 126 | 0 | 7 | 126 |
| Ligands | 3 | 0 | 3 | 3 |
| Avg. *B*-factors (Å$^2$) | | | | |
| Protein | 121.5 | 120.4 | 101.5 | 100.0[1] |
| DNA | 302.9 | — | 73.4 | 100.0[1] |
| Ligand | 68.0 | — | 85.7 | 100.0[1] |
| R.m.s. deviations | | | | |
| Bond lengths (Å) | 0.004 | 0.003 | 0.004 | 0.004 |
| Bond angles (°) | 0.487 | 0.424 | 0.423 | 0.571 |
| Ramachandran plot (%) | | | | |
| Favored | 97.25 | 97.95 | 96.81 | 96.95 |
| Allowed | 2.75 | 2.05 | 3.19 | 3.05 |
| Disallowed | 0.00 | 0.00 | 0.00 | 0.00 |
| MolProbity score | 1.14 | 0.81 | 1.21 | 1.51 |
| Clashscore | 2.22 | 1.00 | 2.33 | 5.96 |
| Poor rotamers (%) | 0.00 | 0.00 | 0.00 | 0.00 |

[1] *B*-factors were manually set and were not refined during structure refinement.

**Supplementary Table S2. Helicase and ATPase rates of FBH1 mutants**

| | Helicase[1] | | ATPase[2] | |
|---|---|---|---|---|
| | $k_{obs}$ (min$^{-1}$) | Rel. to WT | $k_{obs}$ (min$^{-1}$) | Rel. to WT |
| WT | 1.7 ± 0.2 | 1.00 | 517.4 ± 25.9 | 1.00 |
| R447A/K448A | 0.2 ± 0.04 | 0.11 | 552.5 ± 27.6 | 1.07 |
| Q510A | 1.1 ± 0.3 | 0.68 | 552.0 ± 27.6 | 1.07 |
| Δ(507-512) | 0.9 ± 0.1 | 0.52 | 395.4 ± 19.8 | 0.76 |
| D573A/E574A | n.d. | - | 0.5 ± 0.03 | 0.001 |
| H830A | 1.4 ± 0.4 | 0.83 | 574.3 ± 28.7 | 1.11 |
| F836A | 1.5 ± 0.6 | 0.89 | 610.3 ± 30.5 | 1.18 |
| K859A | 0.6 ± 0.1 | 0.34 | 557.6 ± 27.9 | 1.08 |
| C925A/C928A | 0.04 ± 0.04 | 0.02 | 13.9 ± 0.7 | 0.03 |
| Δ(946-end) | 1.8 ± 0.2 | 1.07 | 538.8 ± 26.9 | 1.04 |

[1] Helicase measured with a fork substrate (oligos #8-11, Table S3).

[2] ATPase measured with a splayed arm substrate (oligos #1-2, Table S3).

**Supplementary Table S3. Oligodeoxyribonucleotides used in bulk biochemistry and structural studies**

| Oligo # | Name | Sequence |
|---|---|---|
| **EMSA** | | |
| 1 | FAM40 | (FAM)CTCAGGACTCAGTTCGTCAGCCCTTGACAGCGATGGAAGC |
| 2 | F20.40 | CGAAGGTAGCGACAGTTCCCCTGACGAACTGAGTCCTGAG |
| 3 | FAM40_3'oh_8 | CGCTGTCAAGGGCTGACGAACTGAGTCCTGAG |
| 4 | FAM40_lead2gap | GCTTCCATCGCTCTCAAG |
| 5 | FAM40_lag2gap | GAACTCTCGCTACCTTCG |
| 6 | FAM40_lead8gap | GCTTCCATCGCT |
| 7 | FAM40_lag8gap | TCGCTACCTTCG |
| **Helicase** | | |
| 8 | 54 | ($P^{32}$)ACGCTGCCGAATTCTACCAGTGCCTTGCTCCCTGTAGAAACGGGTGGACGTCCAAGTGGG |
| 9 | 52 | GGGTGAACCTGCAGGTGGGCAAAGATGTCCCAGCAAGGCACTGGTAGAATTCGGCAGCGTC |
| 10 | 56 | CCCACTTGGACGTCCACCCGTTTCTACAG |
| 11 | 53_8gap | TTGCCCACCTGCAGGTTCACCC |
| 12 | 53_8gap_trap | GGGTGAACCTGCAGGTGGGCAA |
| 13 | 9oh_helicase | ($P^{32}$)GGGTGAACCTGCAGGTGGGCAAAGATGTCCCAGC |
| 14 | 53_5gap | TCTTTGCCCACCTGCAGGTTCACCC |
| 15 | 53_5gap_trap | GGGTGAACCTGCAGGTGGGCAAAGA |
| **Fork reversal[1]** | | |
| 16 | 48 | ($P^{32}$)ACGCTGCCGAATTCTACCAGTGCCTTGCTAGGACATCTTTGCCCACCTGCAGGTTCACCC |
| 17 | 50 | GGGTGAACCTGCAGGTGGGCAAAGATGTCC |
| 18 | 50_5gap | GGGTGAACCTGCAGGTGGGCAAAGA |
| 19 | 52 | GGGTGAACCTGCAGGTGGGCAAAGATGTCCCAGCAAGGCACTGGTAGAATTCGGCAGCGTC |
| 20 | 53 | GGACATCTTTGCCCACCTGCAGGTTCACCC |
| 21 | 53_5gap | TCTTTGCCCACCTGCAGGTTCACCC |
| 22 | 53_8gap | TTGCCCACCTGCAGGTTCACCC |
| 23 | A | ($P^{32}$)CGTGACTTGATGTTAACCCTAACCCTAAGATATCGCGTTATCAGAGTGTGAGGATACATGTAG GCAATTGCCACGTGTCTATCAGCTGAAGTTGTTCGCGACGTGCGATCGTCGCTGCGACG |
| 24 | B | CGTCGCAGCGACGATCGCACGTCGCGAACAACTTCAGCTGATAGACACGTGGCAATTGCCTA CATGTATCCTCACACTCTGA |
| 25 | D | CGTCGCAGCGACGATCGCACGTCGCGAACAACTTCAGCTGATAGACACGTGGCAATTGCCTA CATGTATCCTCACACTCTGAATACGCGATATCTTAGGGTTAGGGTTAACATCAAGTCACG |
| **Cryo-EM** | | |
| 26 | EM1-FAM45for_fork | (FAM)CTCAGGACTCAGTTCGTCAGCCCTAGACAGCGATGGAAGCGTCAG |
| 27 | EM2-FAM45lead2gap | CTGACGCTTCCATCGCTGTCTAG |
| 28 | EM3-FAM45rev_fork | GACTGCGAAGGTAGCGACAGATCCCCTGACGAACTGAGTCCTGAG |
| 29 | EM4-FAM45lag8gap | TCGCTACCTTCGCAGTC |

[1] Underlined nucleotides denote mismatches designed to prevent spontaneous fork reversal.

**Supplementary Table S4. DNA substrate construction**

| Figure | Substrate name | Annealed oligos[1] |
|---|---|---|
| 1a | Lead 2, lag 2 gap | 1+2+4+5 |
| 1a | Lead 8, lag 2 gap | 1+2+5+6 |
| 1a | Lead 2, lag 8 gap | 1+2+4+7 |
| 1a | Overhang | 1+3 |
| 1b | Overhang | 13+14 |
| 1b, 5f (bottom) | Fork | 8+9+10+11 |
| 3a | No gap fork | 16+17+19+20 |
| 3b, 5f (top) | Lag gap fork | 16+17+19+21 |
| 3b | Lead gap fork | 16+18+19+20 |
| 3c | Flap | 23+24+25 |
| 5a-e | Cryo-EM | 26+27+28+29 |

[1] Oligo numbers from Table S3.

**Supplementary Table S5. Oligodeoxyribonucleotides used in single-molecule studies**

| Fragment | Oligonucleotide | Sequence |
|---|---|---|
| **Hairpin substrate** | | |
| PCR of dsDNA fragments (connected to DIG and BIO handles) | 453-294 HPLC | GGACAGGCGGCGAGCTGAGGGTTACCGGATAAGGCGCAGCG |
| | 296.R JY0 137 PspOMI-KpnI | ACTTACGCGGGCCCGGTACCGCCAGCATTATGCAGGCCTGGA |
| DIG side adaptor | 295.P-hairpin 5nt over 137A | [Pho]TCAGCTCGCCGCCtgtcccAGCAAGGCACTGGTAG |
| BIO side adaptor | 297.P-hairpin over 137B | [Pho]CCGACTACCAGTGCCTTGCTAGGACAGGCGGCGAGC |
| Short dsDNA hairpin | 250.Loop hairpin | [Pho]TCGGGTCAGATGCCTTTTGGCATCTGAC |
| MT Hairpin DIG handle | 57.FMH_F2_BamHI-ApaI | GCGTAAGTGGATCCGGGCCCCGACTCACTATAGGGAGACCGGC |
| | JOE-R1 | AGTAAGCGCCGTCAGACCAG |
| MT Hairpin BIO handle | 42.FMH_F2_KpnI-PsiI-ScaI | GCGTAAGTGGTACCTTATAAAGTACTCGACTCACTATAGGGAGACCGGC |
| | 209.BsrGI 71short handle | CGATAACCAACTGGCGATG |
| **Fork substrate** | | |
| PCR Hairpin fragment | 248.F-Lambda BsaI 40037 | GCGTAAGTGGTCTCACCGAGCACTACTGGCTGGTTACCAAC |
| | 249.R-Lambda BsaI 41236 | GCTTCCATGGTCTCATTACCACAACTTCCCTGACAAACCG |
| Fork structure | 359.5-Phosphate BsaI | [Pho]GGGCGCATGTATTACTTGGTAGGATCCGTCATAGCTTTAGCGATTTG GGACACTTCATCAAGACTTCCAGAGCAGCCGGAGACATATAGCTACAGG |
| | 293.Template hairpin 30dC | [Pho]GTAACCTGTAGCTATATGTCTCCGCCCCCCCCCCCCCCCCCCCC CCCCCCCCCCTGTGTGTGTGTGTGGTTGTGTGGTGTGTGGTTGTGTGTTG GTGGTTGCATACTTCCGGGAACGCAG |
| Short dsDNA hairpin | 250.Loop hairpin | [Pho]TCGGGTCAGATGCCTTTTGGCATCTGAC |
| dsDNA end compatible with BIO handle | 360.3 BsaI anneal 359 | TGCTCTGGAAGTCTTGATGAAGTGTCCCAAATCGCTAAAGCTATGACGGAT CCTACCAAGTAATACATGC |
| Dig labelling | 253.Primer for Dig | AAAAAAGTGTTGTGTGGTGTTGTTTGGGTGTTGTTTGTGTGTTGTTTGGTGTT GTTTGGGTGTTGTTTGTGTGTTGTTTGCTGCGTTCCCGGAAGTATGC |
| MT Hairpin BIO handle | 361.FMH_F2_BsaI gccc | GCGTAAGTGCCCAGAGACCGGCCTCGAGCCATTTAAG |
| | 209.BsrGI 71short handle | CGATAACCAACTGGCGATG |

### Supplementary Figures

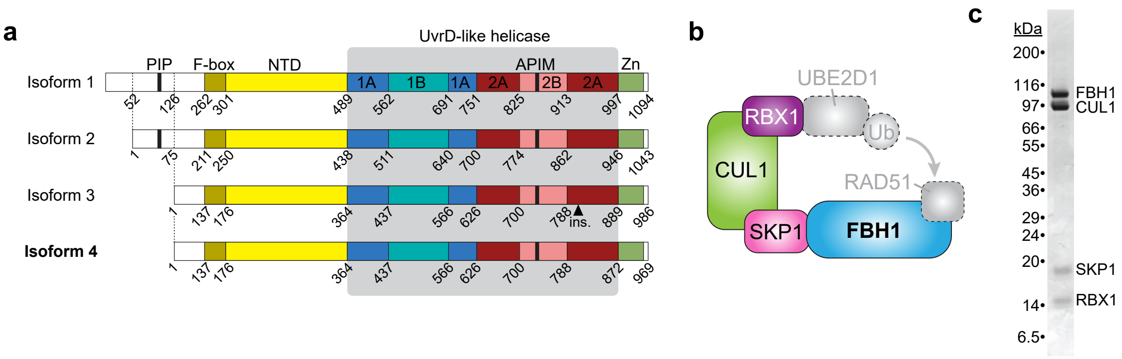


**Figure S1. SCF$^{FBH1}$. a**. Primary structure of FBH1 isoforms 1-4. Residue numbers of isoforms 1 and 4 differ by 125 residues. **b**. Subunit schematic of SCF$^{FBH1}$ (colored) in the context of the E2 ubiquitin ligase (UBE2D1), ubiquitin (Ub), and RAD51 substrates (grey). **c**. Coomassie-stained SDS-PAGE gel of purified SCF$^{FBH1}$ used in this study.

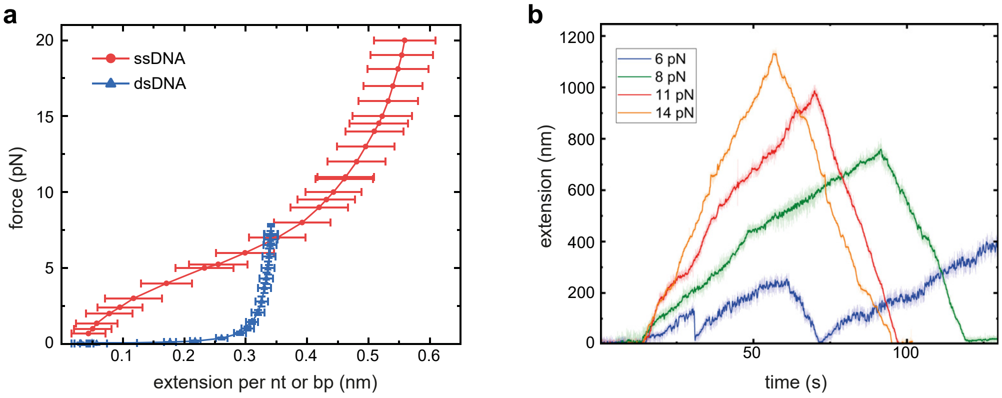


**Figure S2. Force-extension curve allows for the conversion between molecule extension in nm and unwound bp, related to Figure 2. a**. A ssDNA force extension curve in SCF$^{FBH1}$ buffer (mean ± SD, n=25). The curve was obtained using the blocking oligo strategy[7]. **b**. Representative time courses of individual activities of SCF$^{FBH1}$ at different forces before doing the extension conversion from nm to unwound bp (same traces shown in Fig. 2b before extension conversion).

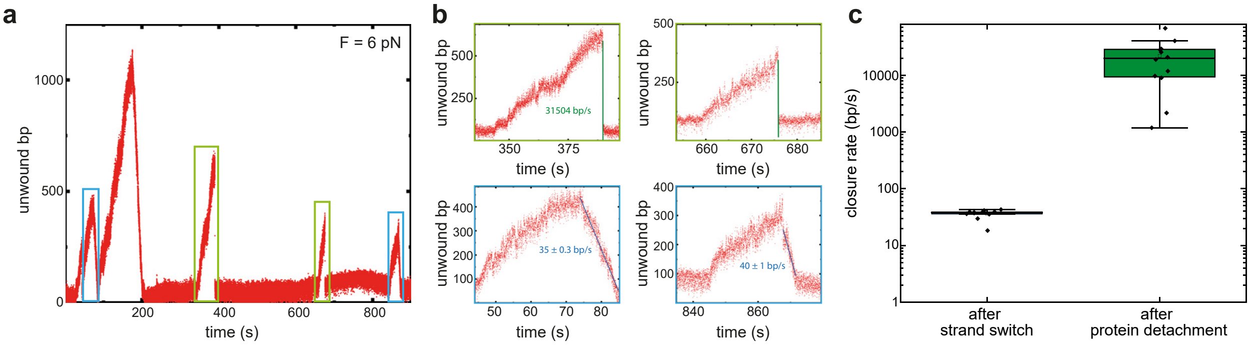


**Figure S3. Hairpin closure after protein detachment is much faster than after strand switching. a**. Representative time-position trace showing both protein-detachment events (green boxes) and strand-switching events (blue boxes). **b**. Zoomed-in views of the highlighted segments from panel a. **c**. Closure-rate distributions for strand-switching events and for spontaneous reclosure following protein detachment. Each box spans the interquartile range (25th–75th percentiles), the internal horizontal line marks the median, and the square symbol denotes the mean. Whiskers extend to the most extreme values within 1.5×IQR of the quartiles.

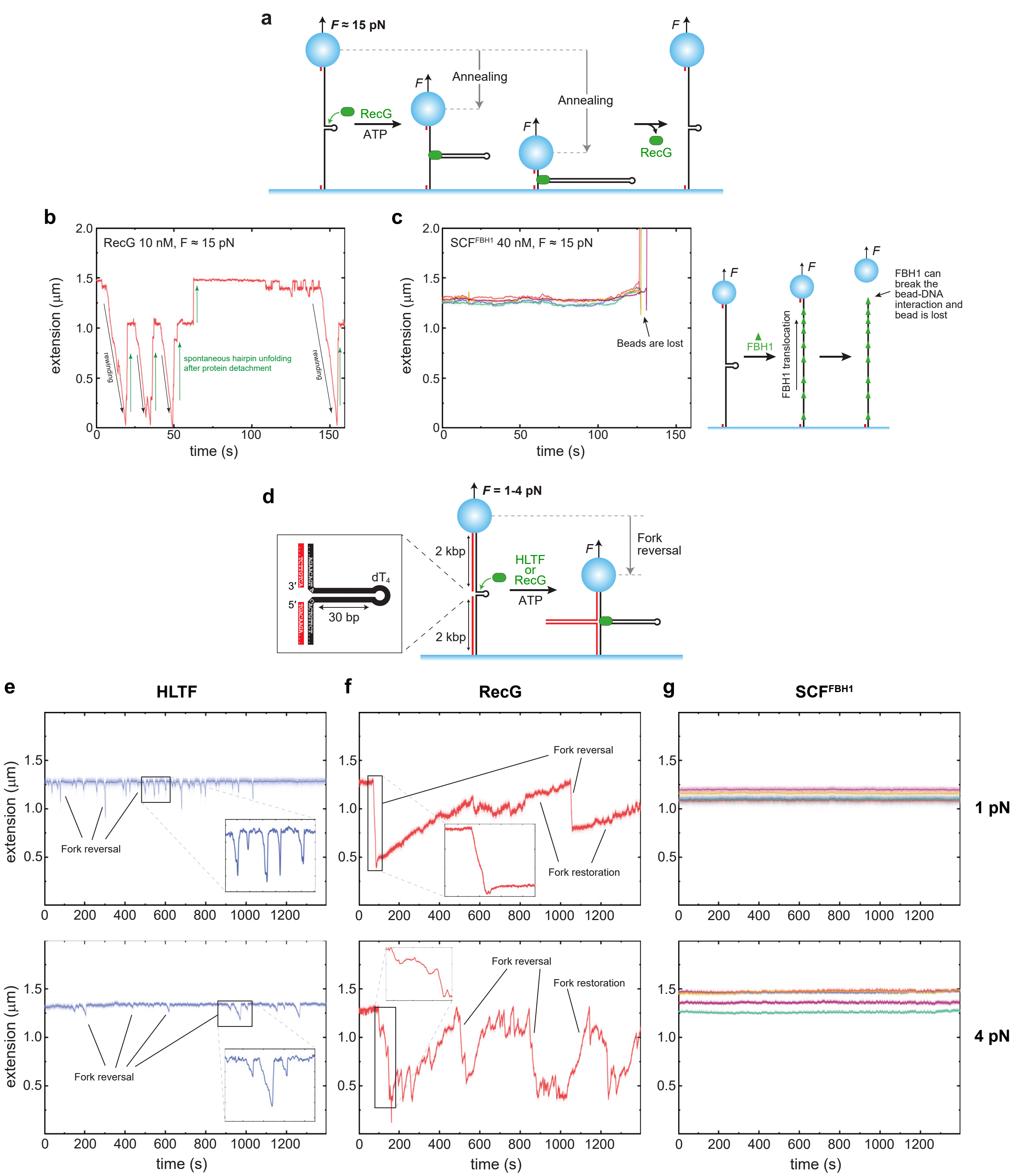


**Figure S4. SCF$^{FBH1}$ lacks annealing activity, related to Figure 3. a**. Schematic of the magnetic tweezers annealing assay. Annealing is detected as a measurable decrease in DNA extension. **b**,**c**. Time-extension traces for 10 nM *Thermotoga maritima* RecG (b) and 40 nM SCF$^{FBH1}$ (c) at ~15 pN opposing force. A schematic explaining the SCF$^{FBH1}$ traces is at right. **d**. Schematic of the magnetic tweezers fork reversal assay. Fork reversal is detected as a measurable decrease in DNA extension. **e-g**. Time-extension traces for 40 nM HLTF (e), 20 nM TmRecG (f), and 40 nM SCF$^{FBH1}$ (g) at 1 pN (top) and 4 pN (bottom) opposing forces. HLTF and RecG reactions contained 1 mM ATP; SCF$^{FBH1}$ reactions contained 2 mM ATP. RecG was purified as described[8]. Colored traces correspond to extension records from individual molecules, measured in parallel.

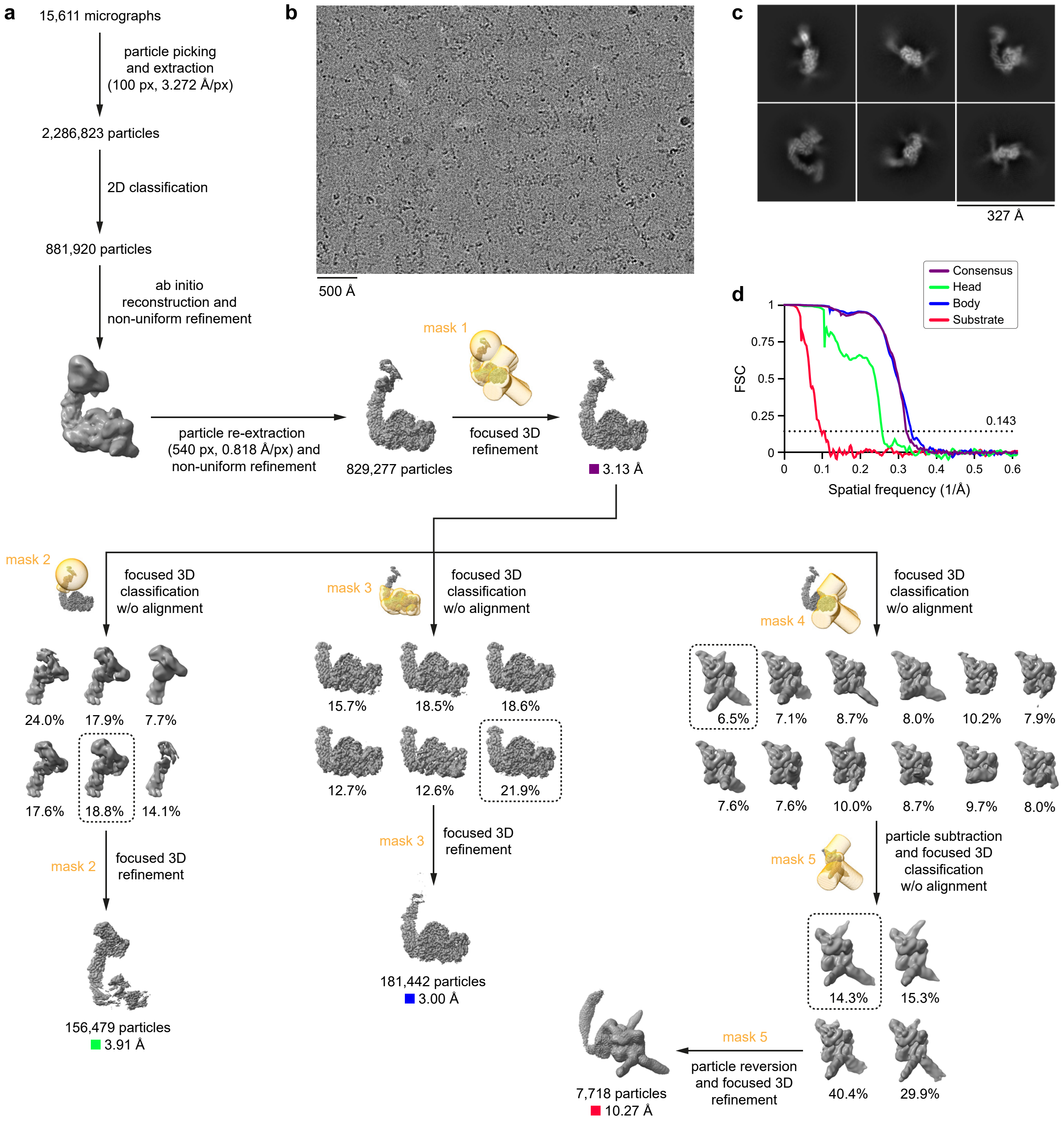


**Figure S5. Cryo-EM reconstruction of the SCF$^{FBH1}$ complex, related to Figures 4, 5, and 7. a.** Data-processing workflow. **b.** Representative micrograph. **c.** Representative 2D class averages. **d.** Gold-standard Fourier shell correlation (FSC) curves calculated from independent half maps. Additional validation metrics are provided in Supplementary Figure S6 and Supplementary Table S1.

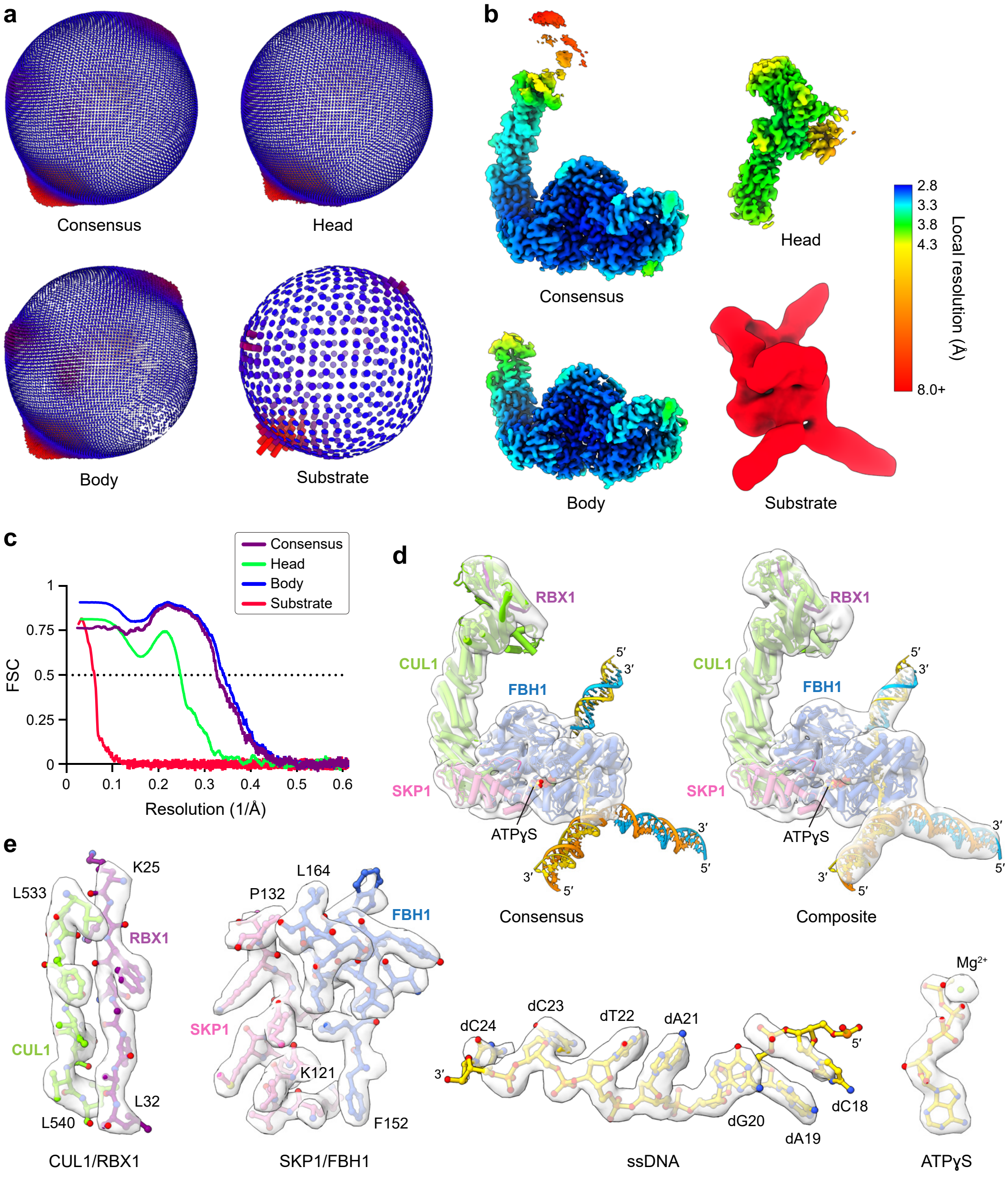


**Figure S6. Structural validation of the SCF$^{FBH1}$ complex, related to Figures 4, 5, and 7. a**. Euler angle distributions for the consensus, head, body, and substrate reconstructions. **b**. Corresponding local resolution estimations. **c**. Model-to-map Fourier shell correlation (FSC) curves calculated using unsharpened full maps. **d**. Model-to-map fits for the complete SCF$^{FBH1}$ complex. Maps were downsampled to a resolution of 10 Å using cryoSPARC. The composite map (right) was generated using PHENIX to combine the consensus, head, body, and substrate maps based upon weighted correlations with the structure of SCF$^{FBH1}$. **e**. Model-to-map fits for selected local regions of the SCF$^{FBH1}$ complex.

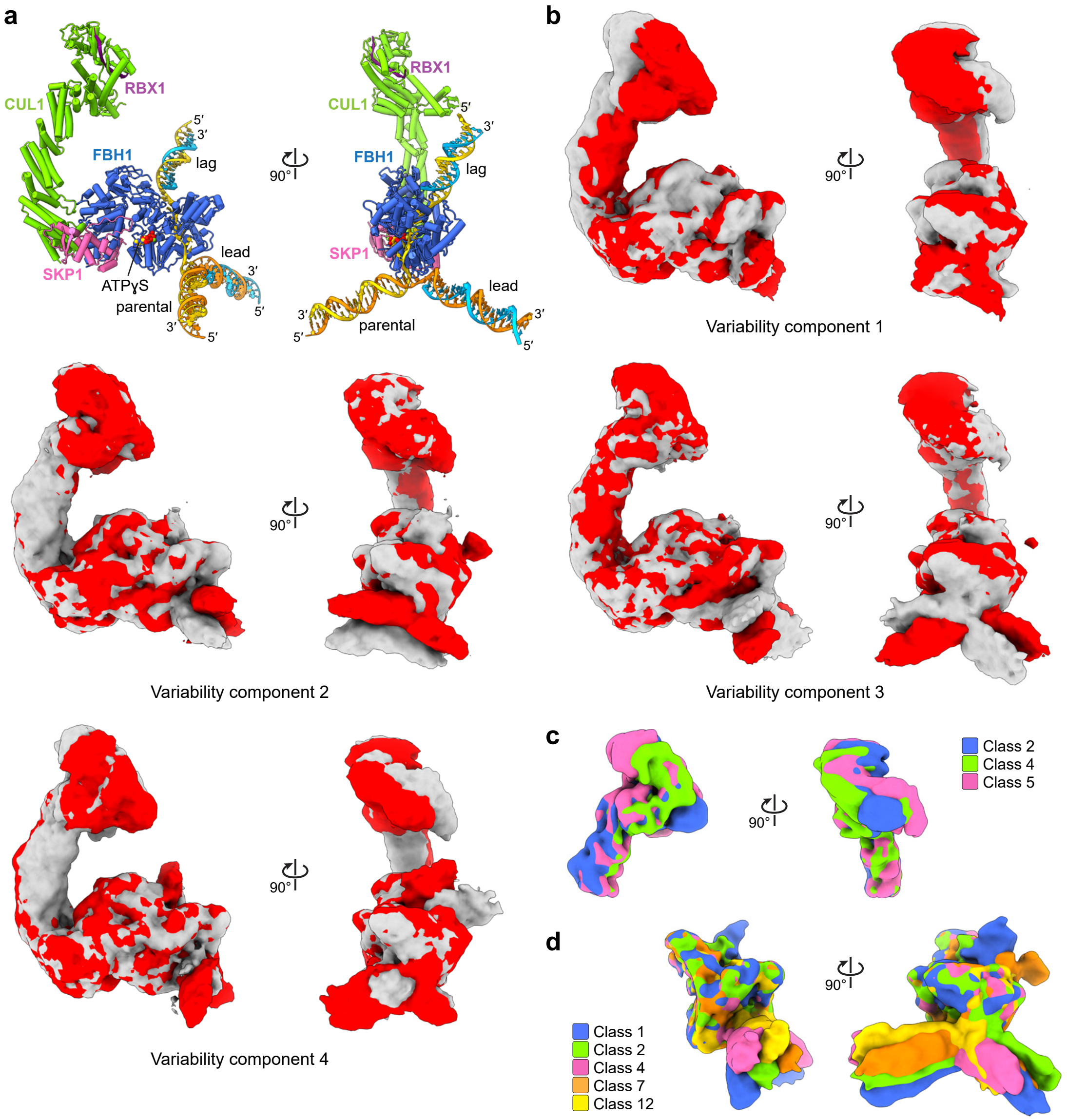


**Figure S7. Characterization of conformational dynamics in the SCF$^{FBH1}$ complex, related to Figures 4, 5, and 7. a**. Orthogonal views of the SCF$^{FBH1}$ structure. The same orientations are shown in panels b and c. **b**. 3D variability analysis of the consensus reconstruction performed using cryoSPARC. Maps were filtered to a resolution of 5 Å. Variability component 1 shows bending of the CUL1 neck and changing interactions in the CUL1/RBX1 head. Variability components 2 and 3 show two distinct motions of the parental and leading DNA duplexes and poorly resolved motion of the lagging duplex. Variability component 4 shows clear motion of the lagging DNA duplex and poorly resolved motion of the parental and leading duplexes. Movies depicting each of the four variability components are provided in the Supplementary Information online. **c**,**d**. Selected 3D classes from the first round of focused 3D classification for the head (c) and substrate (d) reconstructions. Class numbers refer to the 3D classes shown in Supplementary Figure S5a. Head class 5 and substrate class 1 were selected for further processing.

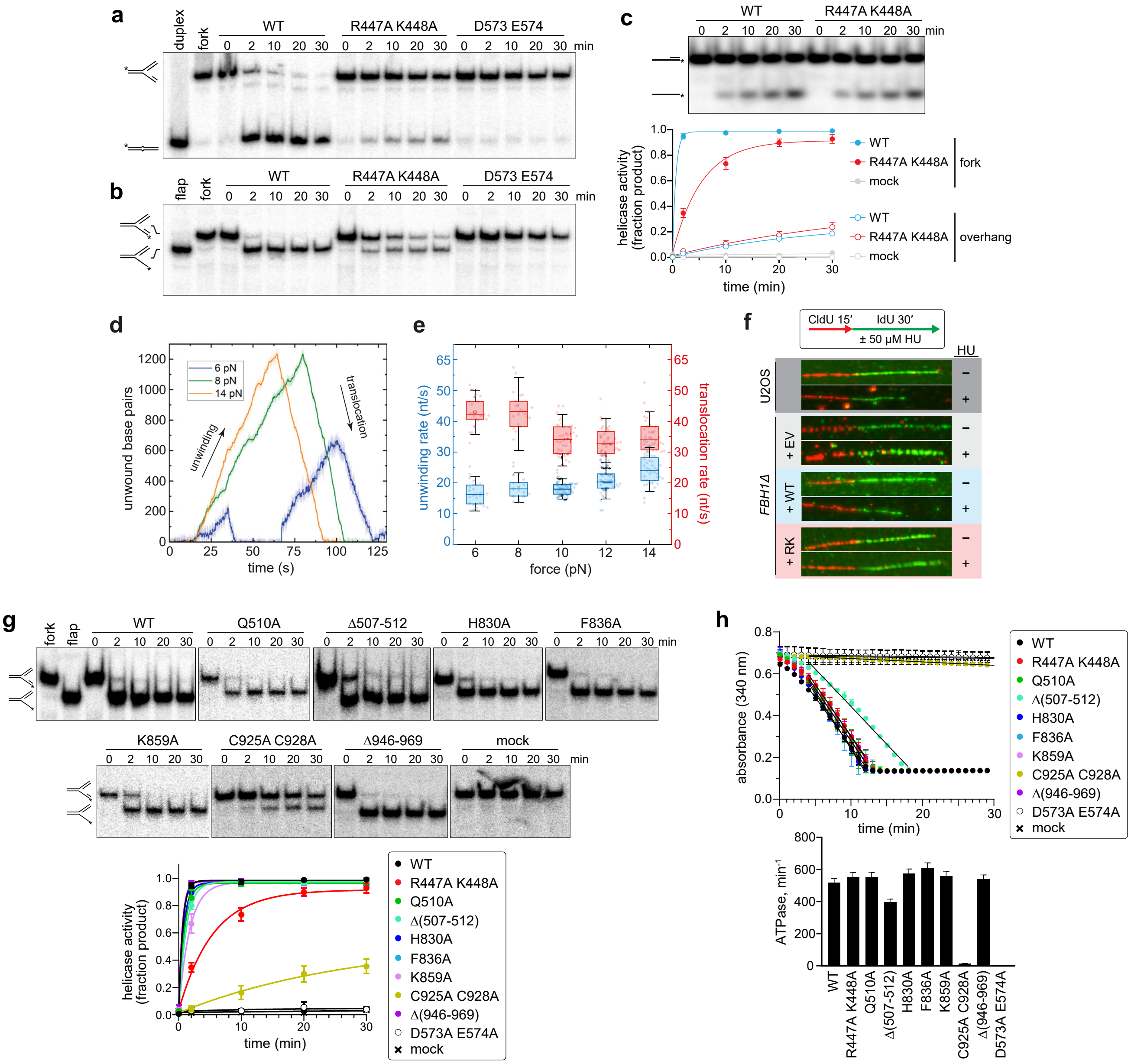


**Figure S8. Fork reversal, helicase, translocation, and ATPase activities of SCF$^{FBH1}$ mutants, related to Figures 5 and 6. a**,**b**. Representative native PAGE separation of fork reversal (a) and helicase (b) substrates and products, quantified in Fig. 5f. **c**. Helicase activity of fork binding mutant for a 3′-overhang substrate. *Top*, representative native PAGE separation of substrate and product. *Bottom*, quantification of data from three independent experiments (mean ± SD, n=3), shown with fork unwinding data from Fig. 5f for comparison. **d**,**e**. Single-molecule magnetic tweezers unwinding and translocation data for R447A K448A at 100 pM. **d**. Representative time courses at different forces. The increase in extension reflects the opening of the hairpin (unwinding activity). The decrease of the extension indicates the closing of the hairpin (translocation activity). **e**. Quantification of duplex unwinding (blue) and ssDNA translocation (red) velocities measured at different forces applied to the hairpin. Box range, 25th-75th percentile; line, median; square, mean; whiskers, 1.5 SD. **f**. Representative DNA combing data, related to Fig. 5g. **g**. Representative native PAGE separation of helicase substrates and products, quantified in Fig. 6f. **h**. ATPase activity, measured using an NADH-coupled regenerative ATPase assay[9]. Top, raw data showing A340 as a function of time (mean ± SD, n=3). Slopes were fit by linear regression and used to determine ATPase rates (bottom). Error bars in the bottom graph denote 5% error. Reactions (100 μL) consisted of 50 nM SCF$^{FBH1}$, 300 nM splayed arm DNA (FAM40+F20.40, Table S3), 40 mM Tris pH 8.0, 50 mM NaCl, 5 mM $MgCl_2$, 1 mM DTT, 1 mM ATP, 3 mM phosphoenolpyruvate, 0.4 mM NADH, 0.7 U pyruvate kinase, and 1 U lactate dehydrogenase, and were carried out at 37°C using a BioTek Synergy H1 plate reader.
